# Hemimetabolous insects elucidate the origin of sexual development via alternative splicing

**DOI:** 10.1101/587964

**Authors:** Judith Wexler, Emily K. Delaney, Xavier Belles, Coby Schal, Ayako Wada-Katsumata, Matthew Amicucci, Artyom Kopp

## Abstract

Insects are the only animals in which sexual differentiation is controlled by sex-specific RNA splicing. The *doublesex* (*dsx*) transcription factor produces distinct male and female protein isoforms (DsxM and DsxF) under the control of the RNA splicing factor *transformer* (*tra*). *tra* itself is also alternatively spliced so that a functional Tra protein is only present in females; thus, DsxM is produced by default, while DsxF expression requires Tra. The sex-specific Dsx isoforms are essential for both male and female sexual differentiation. This pathway is profoundly different from the molecular mechanisms that control sex-specific development in other animal groups. In animals as different as vertebrates, nematodes, and crustaceans, sexual differentiation involves male-specific transcription of *dsx*-related transcription factors that are not alternatively spliced and play no role in female sexual development. To understand how the unique splicing-based mode of sexual differentiation found in insects evolved from a more ancestral transcription-based mechanism, we examined *dsx* and *tra* expression in three basal, hemimetabolous insect orders. We find that functional Tra protein is limited to females in the kissing bug *Rhodnius prolixus* (Hemiptera), but is present in both sexes in the louse *Pediculus humanus* (Phthiraptera) and the cockroach *Blattella germanica* (Blattodea). Although alternatively spliced *dsx* isoforms are seen in all these insects, they are sex-specific in the cockroach and the kissing bug but not in the louse. In *B. germanica*, RNAi experiments show that *dsx* is necessary for male, but not female, sexual differentiation, while *tra* controls female development via a *dsx-*independent pathway. Our results suggest that the distinctive insect mechanism based on the *tra-dsx* splicing cascade evolved in a gradual, mosaic process: sex-specific splicing of *dsx* predates its role in female sexual differentiation, while the role of *tra* in regulating *dsx* splicing and in sexual development more generally predates sex-specific expression of the Tra protein. We present a model where the canonical *tra*-*dsx* axis originated via merger between expanding *dsx* function (from males to both sexes) and narrowing *tra* function (from a general splicing factor to the dedicated regulator of *dsx*).

## INTRODUCTION

Sex determination and sexual differentiation evolve on drastically different time scales. Sex determination (that is, the primary signal directing the embryo to develop as male or female) can be either environmental or genetic. Genetic sex determination can in turn be either male- or female-heterogametic (with or without heteromorphic sex chromosomes), mono- or polygenic, or haplo-diploid, among other mechanisms (1–7). Primary sex determination is among the most rapidly evolving developmental processes. In African clawed frogs, medaka, salmon, and other animals, there are many examples of recently evolved sex-determining genes, so that the primary sex determination genes can differ between closely related species (8–13). The signals initiating male or female development can vary even within species, as seen in cichlids, zebrafish, house flies, and Ranid frogs (5,14–18).

In contrast, sexual differentiation (that is, the set of molecular mechanisms that translates the primary sex-determining signal into sexually dimorphic development of specific organs) tends to remain stable for hundreds of millions of years despite the rapid evolution of both sex determination and the multitude of anatomical, physiological, and behavioral manifestations of sexual dimorphism. This fits the broader “hourglass” pattern of developmental evolution, where the most upstream and downstream tiers of developmental hierarchies diverge more rapidly than the middle tiers (19–21). In the case of vertebrate sexual development, this middle tier is comprised of sex hormones and their receptors, which are largely conserved from mammals to fish, as are the key components of the network of transcription factors and signaling pathways that toggle gonad development between ovary and testis (22–28). In nematodes, key components of the sexual differentiation pathway are conserved between *Caenorhabditis* and *Pristionchus*, which are separated by 200-300 million years (29).

Similarly, the insect sexual differentiation pathway, while unique among metazoans, shows deep conservation across 300 million years of insect evolution (30–37). This pathway has three distinctive features. First, the *doublesex* (*dsx*) transcription factor is spliced into a male-specific isoform in males and a female-specific isoform in females; the two isoforms share a common N-terminal DNA-binding domain but have mutually exclusive C-termini, which lead them to have distinct and often opposite effects on downstream gene expression and morphological development (38–44). Second, alternative splicing of *dsx* is controlled by the RNA splicing factor *transformer* (*tra*); and third, *tra* itself is alternatively spliced so that it produces a functional protein only in females, while in males a premature stop codon results in a truncated, non-functional Tra protein (38–42). Thus, the male-specific *dsx* isoform is produced by default, while the production of the female *dsx* isoform requires active intervention by *tra*. Despite some differences in details, the *transformer*-*doublesex* splicing cascade is conserved among insect orders separated by >300 million years, including Diptera, Coleoptera, and Hymenoptera, indicating that this pathway was already present in the last common ancestor of holometabolous insects (32,33,36,37). The one exception to this rule is found in Lepidoptera, where *tra* has been secondarily lost and *dsx* splicing is regulated instead by a male-specific protein and a female-specific piRNA (38).

Although conserved within insects, the *transformer*-*doublesex* splicing pathway appears to be unique among animals, as nothing similar has been observed in any other animal group. The differences in the molecular basis of sexual differentiation between insects, nematodes, and vertebrates are fundamental. In insects, *dsx* and *tra* operate in a mostly cell-autonomous manner, as evidenced for example by dramatic insect gynandromorphs (38–41). In vertebrates, sexual differentiation is largely non-cell-autonomous. During embryonic development, the originally bipotential vertebrate gonad is biased to become either ovary or testis under the influence of the primary sex-determining signal, which can be either genetic or environmental (22,38–40,40). Subsequently, male or female hormones produced by the gonad act through nuclear receptor signaling to direct somatic tissues to differentiate in sex-specific ways. In rhabditid nematodes such as *Caenorhabditis elegans*, a multi-step cell signaling cascade involving secreted ligands, transmembrane receptors, and post-translational protein modification results in sex-specific expression of several transcription factors that direct male or female/hermaphrodite development of particular cell lineages (38–41).

The deep conservation of fundamentally different mechanisms of sexual differentiation in different animal phyla presents an intriguing question: how do such disparate mechanisms evolve in the first place? Comparison of nematode, crustacean, insect, and vertebrate modes of sexual differentiation suggests that the insect-specific mechanism based on alternative splicing of *dsx* evolved from a more ancient mechanism based on male-specific transcription of an ancestral *dsx* homolog. *dsx* is part of the larger Doublesex Mab-3 Related Transcription factor (DMRT) gene family, which is apparently the only shared element of sexual differentiation pathways across metazoans (42,43). *Dmrt1* is involved in the specification and maintenance of testis fate in mammals and other vertebrates, while repressing ovarian differentiation (26,44,45). Due to their function in the androgen-producing Sertoli cells, vertebrate *Dmrt1* genes also play a crucial role in the development of secondary sexual characters. In *C. elegans*, *mab-3* and several other DMRT family genes are responsible for the specification of various male-specific cells, structures, and behaviors (46–50). However, in both vertebrates and nematodes, the DMRT genes involved in sexual differentiation are not spliced sex-specifically, but are transcribed in a predominantly male-specific fashion, promote the development of male-specific traits, and are dispensable for female sexual differentiation. In contrast, the insect *dsx* acts as a bimodal switch that plays active roles in both male- and female-specific differentiation through its distinct male and female splicing isoforms. In the absence of Dsx, both males and females develop as intersexes with phenotypes intermediate between males and females (51,52).

In the closest relative of insects that has been studied to date, the branchiopod crustacean *Daphnia magna*, *dsx* acts in a manner similar to vertebrates and nematodes rather than insects: it is transcribed male-specifically, is not alternatively spliced, controls the development of male-specific structures, and is dispensable in females (53). The *D. magna tra* gene is not spliced sex-specifically, does not differ in expression between males and females, and does not regulate *dsx* (Kato et al, 2010). *dsx* also shows male-biased transcription and no sex-specific splicing in the shrimp *Fenneropenaeus chinensis* (54) and in the mite *Metaseiulus occidentalis* (54). Thus, the origin of the *transformer-doublesex* splicing cascade was a key event in the evolution of sexual differentiation in insects.

To understand the evolutionary transition from a transcription-based to a splicing-based mode of sexual differentiation, it is necessary to focus on the phylogenetic interval between branchiopod crustaceans and holometabolous insects. This interval spans several crustacean groups, non-insect Hexapods, and many hemimetabolous insect orders, and encompasses a broad range of body plans and developmental mechanisms. Unfortunately, the development of these groups remains poorly studied compared to holometabolous insects; in particular, very little is known about sexual differentiation in hemimetabolous insects.

To explore the origin of the insect sexual differentiation pathway based on the *transformer-doublesex* axis, we examined the expression of these genes in three hemimetabolous insects: the kissing bug *Rhodnius prolixus* (Hemiptera), the louse *Pediculus humanus* (Phthiraptera), and the German cockroach *Blattella germanica* (Blattodea). We find that distinct male and female *dsx* isoforms are conserved through Blattodea, the most basal insect order studied to date. However, only *R. prolixus* shows the canonical sex-specific pattern of *tra* splicing. In *B. germanica*, we show that *tra* nevertheless controls *dsx* splicing and is necessary for female but not male sexual development, as in holometabolous insects. Surprisingly, the *B. germanica dsx* gene is required for male sexual differentiation, but appears to be dispensable in females; in this respect, the cockroach is more similar to crustaceans and non-arthropod animals than to holometabolous insects. Together, our results suggest that the splicing-based mode of sexual differentiation based on the *transformer*-*doublesex* regulatory pathway has evolved in a gradual, mosaic fashion in hemimetabolous insects.

## RESULTS

### *doublesex* and *transformer* orthologs are present in hemimetabolous insects

Although *dsx*, *Dmrt1*, and other DMRT genes have prominent roles in sexual differentiation (22,42,43,46,55), many DMRT paralogs are involved in other developmental processes, and most have phylogenetically restricted distributions (56). The number of DMRT paralogs varies among animal taxa; for example, *D. melanogaster* has four, while *C. elegans* has 11 (56). To identify *dsx* orthologs in hemimetabolous insects for functional study, we performed a phylogenetic analysis of arthropod DMRT genes (Supplemental Table 1). By mining arthropod gene models, we recovered the deeply conserved Dmrt11E, Dmrt93B, and Dmrt99B subfamilies in addition to a large clade containing *dsx* orthologs. The *dsx* clade contained sequences from several basal insect orders, crustaceans, and chelicerates (Supplemental Figure 1), and included experimentally characterized *dsx* genes from the branchiopod crustacean *D. magna* (NCBI accession number BAJ78307.1), the decapod crustacean *F. chinensis* (AUT13216.1), and the mite *M. occidentalis* (XP003740429.2 and XP003740430.1). In these species, the *dsx* genes are not spliced sex-specifically, but are transcribed in a strongly male-biased fashion (53,54,57). *dsx* genes from holometabolous insects, which direct male and female differentiation via alternative spliceforms, are also present in this clade. There are marked departures in this gene tree from the arthropod species tree. Notably, many hemipteran *dsx* sequences group with chelicerates and crustaceans instead of insects, although with low support. The clustering of crustacean and chelicerate *dsx* genes, which control male sexual development via male-specific upregulation, with holometabolous *dsx* genes that control male and female development via alternative spliceforms indicates that sex-specific *dsx* isoforms evolved after the origin of the *dsx* clade.

Puzzled by the clustering of hemipteran *dsx* genes with those from chelicerates and crustaceans, we conducted a synteny analysis. While the robustness of this analysis is dependent on the quality of genome assembly, the transcription factor *prospero (pros)* is on the same scaffold as *dsx,* with an intervening distance between 17 and 245 kb (Supplemental Table 2), in seven different holometabolous and hemimetabolous insect orders (Ephemeroptera, Blattodea, Phthiraptera, Thysanoptera, Hymenoptera, Coleoptera and Lepidoptera). However, in Hemiptera, tBLASTn searches of the *prospero*-containing scaffolds, ranging in size from ∼120 kb to 17 Mb (mean 2.75 Mb), did not identify neighboring DMRT genes (Supplemental Table 3). Recent work in the planthopper *Nilaparvata lugens* suggests that at least some of the Hemipteran *dsx* genes are alternatively spliced and are necessary for male development (58). We conclude that hemipterans have *dsx* orthologs, but the synteny between *pros* and *dsx* was lost in the common ancestor to true bugs, and Dsx protein sequences evolved rapidly in this clade.

Transformer proteins have repetitive and simple sequences that evolve rapidly, posing greater difficulties for phylogenetic analysis. Previous studies of insect *tra* genes have identified these genes via the characteristic arginine/serine rich (RS) domain (33), used BLAST to detect a *tra* candidate and demonstrated congruence between a species tree and a gene tree (53), or simply concluded that a candidate *tra* gene was indeed *transformer* after knocking it down and obtaining a sex-specific phenotype (36). *tra* is related to, but is not part of, a large family of genes structurally defined by the presence of an RNA binding domain and at least one RS domain (59). These proteins, called SR family genes for their amino acid composition, commonly function in pre-mRNA splicing (59). Because most insect *tra* genes lack an RNA binding domain (see below), they are classified as RS-like proteins. Our maximum likelihood phylogenetic analysis of Tra proteins, along with three previously studied SR family genes (Transformer-2, which is not an ortholog of Transformer; SFRS; and Pinin) shows that putative *tra* genes from some hemimetabolous insects including *R. prolixus, P. humanus*, and *B. germanica* cluster with experimentally characterized holometabolous *tra* genes and with the *Daphnia tra*, although with low support (Supplemental Figure 2). Despite poor phylogenetic resolution, the putative *tra* genes identified in *R. prolixus*, *P. humanus,* and *B. germanica* provided an avenue for experimental analysis of sexual differentiation in hemimetabolous insects.

Prior work has shown that *dsx* is arthropod-specific (56); the present analyses confirm that *dsx* was probably present in the last common ancestor of arthropods. More research is needed to pinpoint the origin of *tra*, but the poorly conserved and highly disorganized nature of Tra proteins suggests that this problem might be intractable.

### *tra* splicing follows the holometabolous sex-specific pattern in *R. prolixus* but not in *P. humanus* or *B. germanica*

Insect Tra proteins are defined by three key features: (1) an arginine-serine rich (RS) domain, (2) a proline rich domain, and, (3) for all non-*Drosophila tra* sequences, an auto-regulatory CAM domain named for the three genera in which it was discovered, *Ceratitis*-*Apis*-*Musca* (60–62). In the holometabolous insect orders Diptera, Coleoptera, and Hymenoptera, a premature male-specific stop codon truncates the *tra* coding sequence, making the male Tra protein unable to regulate *dsx* splicing (33,36,63–65). These insects produce a male-specific *dsx* isoform by default, while females produce a female-specific *dsx* isoform as a result of having functional Tra. To understand when *tra*-dependent alternative splicing of *dsx* evolved, we identified *tra* isoforms in males and females from three hemimetabolous insect orders (Hemiptera: *R. prolixus*; Phthiraptera: *P. humanus*; and Blattodea: *B. germanica*). We find that the characteristic holometabolous-like pattern of sex-specific splicing of *tra*, with a male-specific premature stop codon, is present in *R. prolixus* but not in *P. humanus* or *B. germanica*. Instead, *P. humanus* and *B. germanica* display novel patterns of alternative splicing, which include the presence of female-biased truncated *tra* isoforms and an interrupted CAM domain.

#### Rhodnius prolixus

In *R. prolixus*, we identified a tandem duplication of *tra* on scaffold KQ034193 (Rhodnius-prolixus-CDC_SCAFFOLDS_RproC3, v.1) via tBLASTn searches of the *R. prolixus* genome (66). In the phylogenetic tree of arthropod SR proteins (Supplemental Figure 2), the two *R. prolixus* paralogs (*RpTraA* and *RpTraB*) cluster within a clade that also contains Dipteran, Hymenopteran, and Coleopteran Tra proteins. The downstream *R. prolixus tra* paralog, which we call *RpTraB*, encodes conserved Tra amino acid residues at its N-terminus, including the CAM domain and an RS domain, but is truncated at the C-terminus and lacks a proline-rich domain (Supplemental Figure 3). In contrast, the upstream *RpTraA* paralog encodes a full-length Tra protein with a C-terminal proline-rich domain. Interestingly, *RpTraA* is also predicted to contain a partial RNA Recognition Motif (RRM) (Supplemental Figure 4), a feature which has not been described in previous studies of Tra proteins. However, when we inspected predicted protein domains in Tra sequences from Branchiopoda, Copepoda, Blattodea, Hemiptera, Hymenoptera, Coleoptera, and Diptera with CCD/SPARCLE software via NCBI (67), we found putative RRM domains in *Apis mellifera* CSD, *R. prolixus* TraA, a predicted Tra protein from the copepod crustacean *Tigriopus californicus*, and the cockroach *B. germanica* Tra ortholog (Supplemental Figure 4). The absence of the RRM domain from most arthropod Tra sequences suggests that insect Tra proteins once had a functional RRM domain but lost RNA-binding activity over time, perhaps due to the association between Tra and the RNA-binding protein Tra-2 (68). *RpTraA* and *RpTraB* are the first reported *tra* duplicates outside of Hymenoptera, where *tra* duplications sometimes result in functional innovation (33,69).

To investigate whether *RpTraA* is expressed similarly to holometabolous *tra* orthologs, we conducted 5’ and 3’ Rapid Amplification of cDNA Ends (RACE) to obtain the sets of isoforms expressed in males and females. We also generated *de novo* transcriptome assemblies using Illumina sequencing and Trinity software (70). We were unable to recover *RpTraB* from adult male or female gonad transcriptomes, nor could we recover any *RpTraB* product by RT-PCR. We found two female-specific *RpTraA* transcripts (*RpTraA_3* and *RpTraA_4*) and two male-specific transcripts (*RpTraA_1* and *RpTraA_2*) (Figure 1A). All transcripts were identified as sex-specific based on their presence/absence in male and female RACE and RNA-seq libraries, and their sex-specificity was further confirmed by RT-PCR (Figure 1B). As in the holometabolous *tra* splicing, both male-specific transcripts (*RpTraA_1* and *RpTraA_2*) contain stop codons near the N-terminus and thus encode truncated proteins that are almost certainly incapable of regulating RNA splicing (Figure 1A). Only the *RpTraA_4* transcript encodes a complete Tra protein with the CAM domain followed by an RS domain, RRM domain, and a proline-rich domain (Figure 1A). The female-specific *RpTraA_3* lacks the stop codon found in male *RpTraA* transcripts, but has truncated RS and CAM domains and lacks the proline-rich domain. This pattern – a male-specific premature stop codon that is located in an alternatively spliced exon and results in the production of a functional Tra protein in females but not in males – is identical to the pattern seen in holometabolous insects including *D. melanogaster*, *A. mellifera*, and *Tribolium castaneum* (33,36,64). Our detection of truncated Tra isoforms in males but not females in a Hemipteran suggests that at least some elements of the sexual differentiation mechanism based on the alternative splicing of *tra* may predate holometabolous insects.

**Figure 1:**
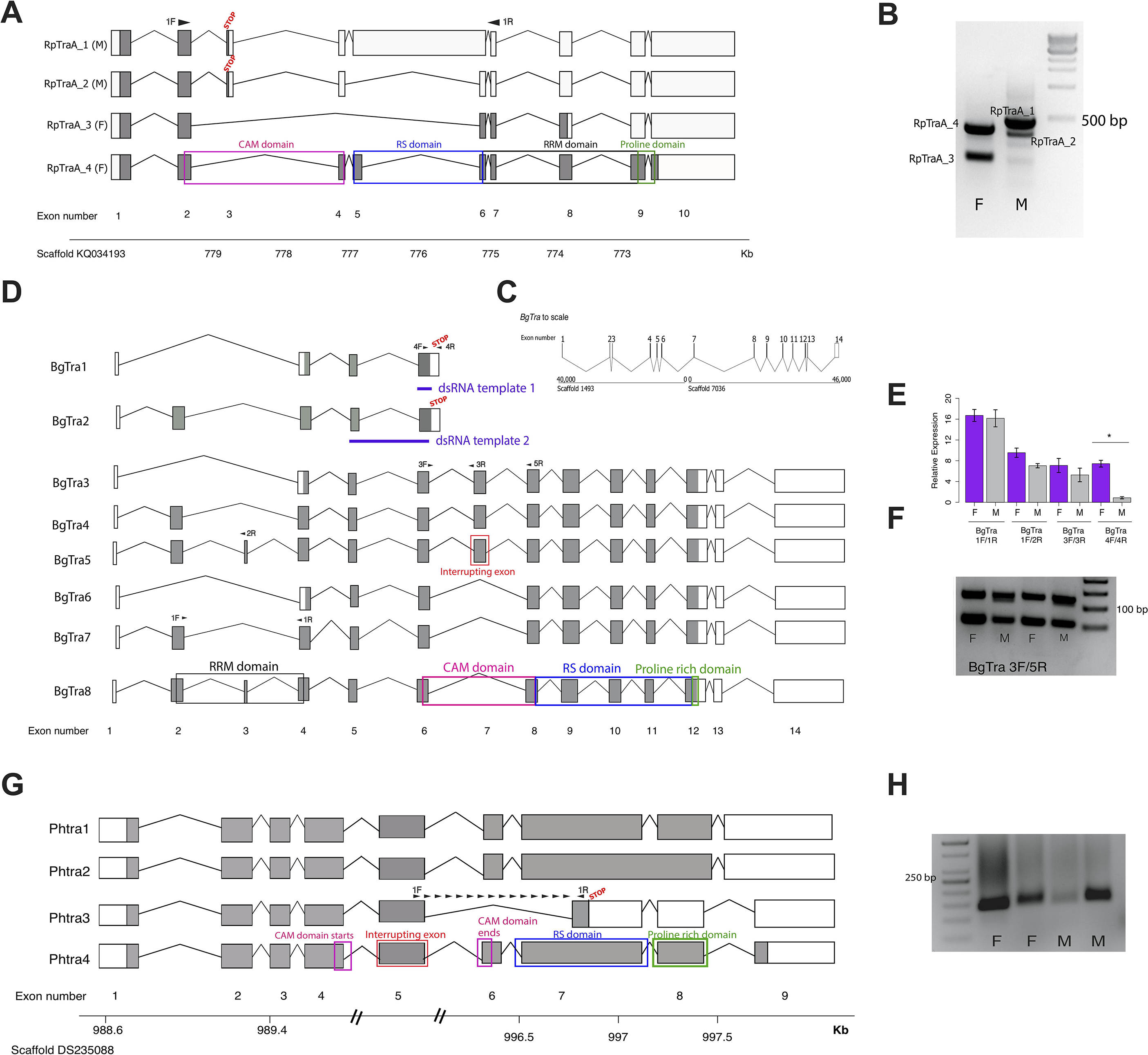
Transformer splicing follows the holometabolous pattern in the kissing bug *Rhodnius prolixus*, but not in the cockroach *Blattella germanica* or louse *Pediculus humanus*. Schematics showing transformer transcripts isolated from *R. prolixus*, *B. germanica*, and *P. humanus*. Coding sequences are in gray, UTRs are in white. (A) Four transcripts isolated from *RpTraA* with 5’ and 3’ RACE and confirmed by RNA-seq. Sex-specificity of each transcript is noted in parentheses at the end of the transcript name. Stop codon in the male-specific 3rd exon is indicated with “STOP.” Predicted Transformer protein domains are shown with colored boxes and labels, using *RpTraA_4* as example. (B) PCR on male and female cDNA with *RpTraA* primers (RpTra_checkF +RpTra_checkR in Supplemental Table 5) indicated by black arrows in (A) confirms the sex-specificity of each transcript. (C) *BgTra* spans two scaffolds in the *B. germanica* genome. Genome coordinates are shown at the beginning and end of each scaffold. Both scaffolds continue beyond *BgTra*; scaffold 1493 is 552 kb and scaffold 7036 is 55 kb. (D) Alternative inclusion of exons 2, 3, 7, and intron read-through after exon 6 result in eight BgTra transcripts. Nucleotides which code for important protein domains are color-coded as in panel A, using *BgTra8* and *BgTra5* as examples. In *BgTra1* and *BgTra2*, exon 6 extends ∼500 bp further 3’ than in all other transcripts. The 3’ end of exon 6 in *BgTra3-8* falls six bp upstream of the stop codon truncating the conceptual translation of *BgTra1* and *BgTra2*. Exon 7 codes for 47 amino acids that interrupt the CAM domain, which spans the end of exon 6 and start of exon 8. Purple lines show the locations of dsRNA used in functional experiments. Black arrows indicate locations of primers used for RT-PCR and qPCR. (E) qPCR expression of different *BgTra* isoforms. The truncated transcripts *BgTra1* and *BgTra2* are female-biased, while the remaining transcripts are sexually monomorphic. (F) RT-PCR showing that both males and females express transcripts with and without the exon interrupting the CAM domain. The shorter band contains *BgTra* isoforms that skip exon 7 and contain an intact CAM domain; the larger product contains exons 6, 7, and 8. (G) Four *PhTra* transcripts isolated from *P. humanus* by 5’ and 3’ RACE. *PhTra3* has a premature stop codon and is found in both sexes, as shown by RT-PCR on male and female cDNA in panel (H). Forward primer for PCR shown in (H) spanned the exon 5/7 junction (marked with black arrows in panel (G)); reverse primer for this PCR is in exon 7 (also noted with a black arrow). All *PhTra* isoforms contain the “interrupting exon” inside of the CAM domain. Double bars on scaffold in panel (G) represent breaks in scaffold scale.

#### Blattella germanica

To characterize *tra* isoforms in male and female German cockroaches, we sequenced full-length transcripts using a combination of PacBio RNA Isosequencing and 5’ and 3’ RACE on male and female samples. Our tBLASTn search of the *B. germanica* genome revealed one putative *tra* ortholog (*BgTra*) based on its clustering with *Apis* and *Rhodnius tra* genes (Supplemental Figure 2). Both PacBio RNA Isosequencing and 5’/3’ RACE revealed that this single *BgTra* gene spans over 86 kb of genomic sequence on two separate scaffolds (genome assembly: i5K Bger Scaffolds v.1) (Figure 1C). The gene model of *BgTra* contains all previously described functional domains of insect Tra proteins – a CAM domain, an RS domain, and a proline rich domain (Figure 1D). We discovered eight *BgTra* transcripts that vary in their inclusion of these domains. The first class of transcripts (*BgTra 1* and *2*), stops in the middle of the CAM domain and lacks the RS and proline rich domains. The second class (*BgTra 6-8*) has an intact CAM domain, while in the third class (*BgTra 3-5*) the CAM domain is interrupted by an extra exon (exon 7 in Figure 1D; Supplemental Figure 5). In addition, variable inclusion of exons 2 and 3 affects the predicted RRM domain in all three classes of *BgTra* transcripts.

Unlike holometabolous insects and *R. prolixus*, we did not find a premature male-specific stop codon in any *tra* transcripts. Surprisingly, the short isoforms (class 1: *BgTra1* and *BgTra2*) with a truncated CAM domain and lacking the RS and proline rich domains had strongly female-biased expression (Figure 1E). RT-PCR and qPCR show that all other isoforms (*BgTra 3-8*) are sexually monomorphic (Figures 1E, 1F). PCR using primers that span the CAM domain amplify the “uninterrupted” CAM domain as well as the “interrupted” CAM domain in both males and females (Figure 1F). This pattern is distinct and unusual – in other insects, sex-specific, prematurely truncated *tra* transcripts are invariably male-biased (60).

#### Pediculus humanus

The louse *P. humanus* shows a pattern of *tra* splicing similar to that in the cockroach. Using a tBLASTn search followed by 5’ RACE, we identified a single gene, *PhTra,* in the *P. humanus* genome, which clustered with other *tra* orthologs in our phylogenetic analyses (Supplemental Figure 2). Using 5’ and 3’ RACE and Illumina RNA-seq followed by de novo transcriptome assembly, we identified four *PhTra* isoforms (Figure 1G). *PhTra1* and *PhTra4* were found in both male and female RACE libraries, as well as in the female RNA-seq assembly. *PhTra2* and *PhTra3* were amplified in only the male RACE library, and were not found in either male or female transcriptome assemblies. *PhTra3* differs from the other isoforms in containing a premature stop codon in the middle of the RS domain (Figure 1G). RT-PCR shows that this isoform is present in both males and females (Figure 1H). The remaining three *PhTra* isoforms encode full-length proteins including an RS-domain and a proline-rich domain, but contain an exon that interrupts the CAM domain, as in some *BgTra* isoforms (Figure 1D, Supplemental Figure 5). The location of this exon is conserved between *PhTra* and *BgTra*. In both *tra* orthologs, the CAM domain is interrupted immediately before two highly conserved resides – a glutamic acid followed by a glycine. Remnants of this interrupting exon, which is only found in hemimetabolous *tra* orthologs, are detected in a conserved exon junction right before these two amino acids in the *tra* orthologs of holometabolous insects (61).

Overall, our comparison of *R. prolixus, B. germanica*, and *P. humanus* shows that the sex-specific splicing of *tra*, which is typical of holometabolous insects and limits Tra protein function to females, is found in some but not all hemimetabolous insects.

### *doublesex* splicing follows the holometabolous, sex-specific pattern in some but not all hemimetabolous insects

In holometabolous insects, the Doublesex transcription factor has male- and female-specific isoforms. The male and female proteins share the DNA-binding DM domain and an oligomerization domain that promotes homodimerization of the transcription factor. This domain is essential to Dsx function, as the protein can only bind DNA efficiently as a dimer (71,72). Male- and female-specific exons at the 3’ end of *dsx* transcripts are responsible for sex-specific expression of Dsx target genes. We recovered one *dsx* ortholog each from *R. prolixus* (*RpDsx*), *P. humanus* (*PhDsx*), and *B. germanica* (*BgDsx*) (Supplemental Figure 1). To test for the presence of sex-specific *dsx* isoforms, we isolated *dsx* transcripts using PacBio Isosequencing, Illumina RNA-sequencing, and 5’ and 3’ RACE. In all three species, we identified alternatively spliced *dsx* transcripts that follow the canonical holometabolous pattern characterized by a common N-terminus and mutually exclusive C-termini. Sex-specific *dsx* splicing is conserved through *B. germanica*, although in *P. humanus* both alternatively spliced isoforms are sexually monomorphic.

#### Rhodnius prolixus

5’ and 3’ RACE revealed three *RpDsx* isoforms, one of which was isolated from female RNA samples (*RpDsx1*) and two from male samples (*RpDsx2* and *RpDsx3*) (Figure 2A). We also retrieved *RpDsx1* from our Trinity *de novo* transcriptome of *R. prolixus* female gonads; we were unable to retrieve any *dsx* isoforms from our male *R. prolixus* transcriptome. RT-PCR confirmed that *RpDsx1* was indeed female-specific, and qPCR showed female-specific expression of *RpDsx1* and male-specific expression of *RpDsx2* and *RpDsx3* (Figure 2B). *RpDsx1* and *RpDsx2* had low expression in *R. prolixus* gonads, whereas *RpDsx3* was expressed at a 6- to 8-fold higher level than either *RpDsx1* or *RpDsx2* (Figure 2B).

**Figure 2:**
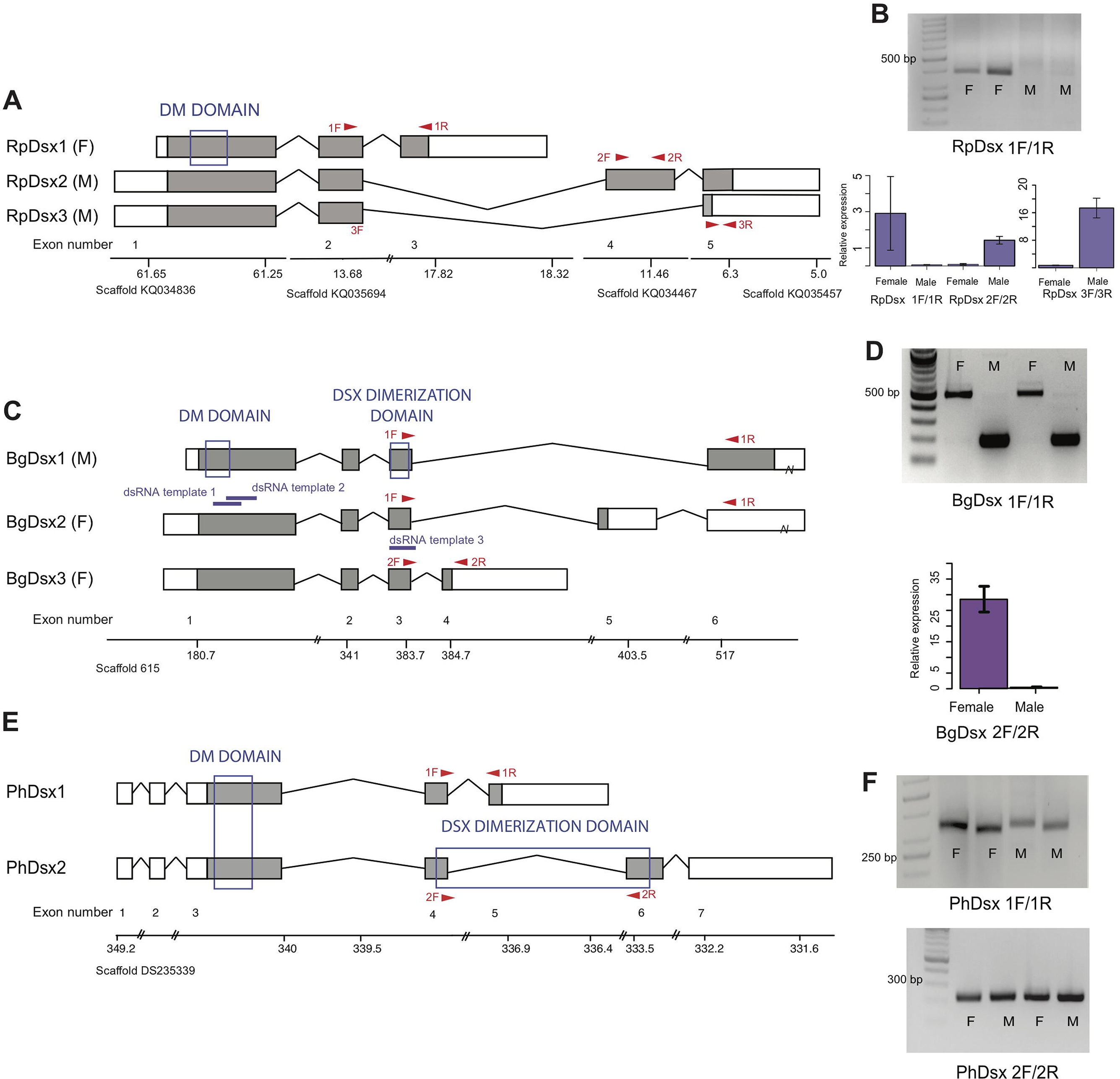
*doublesex* shows sex-specific splicing in some but not all hemimetabolous insects. Schematics showing *doublesex* transcripts isolated from *Rhodnius prolixus*, *Blattella germanica*, and *Pediculus humanus*. Coding sequences are in gray and UTRs in white. Sex-specificity of each transcript is noted in parentheses at the end of the transcript name. Transcripts are shown mapped to genomic scaffolds; double bars show breaks in scale. Functional domains are labeled and indicated by blue boxes. (A) Three *dsx* transcripts isolated from *R. prolixus* span multiple genomic scaffolds. *RpDsx1* was isolated from 3’ and 5’ RACE libraries made from female RNA; *RpDsx2* and *RpDsx3* were cloned from RACE libraries synthesized from male RNA. (B) Top panel: RT-PCR showing female-specificity of *RpDsx1*, using primers in exons 2 and 3, as indicated with red arrows in (A). Bottom panel: qPCR showing female-specific expression of *RpDsx1*, and male-specific expression of *RpDsx2* and *RpDsx3,* using primers indicated with red arrows in (A). Forward primer 3F spanned the exon 2/5 junction. (C) Three *dsx* transcripts isolated by 3’ and 5’ RACE and PacBio Isosequencing from *B. germanica*. The 3’ UTR of *BgDsx1* and *BgDsx2* contains a retrotransposon (marked with a break) and cannot be mapped to a single scaffold of the roach genome. Purple lines show the locations of dsRNA used for RNAi. Red arrows show primer locations for RT-PCR and qPCR. (D) Sex-specific expression of *BgDsx* isoforms. Top panel, PCR on cDNA of adult male and female fat body and reproductive tract with primers in exons 3 and 6. The larger band found exclusively in females contains the alternatively included exon 5, as confirmed by Sanger sequencing. Bottom panel, qPCR for exon 3/4 junction shows amplification only in females. (E) Two *dsx* transcripts isolated by 3’ and 5’ RACE from *P. humanus* male and female samples. (F) RT-PCR shows the presence of both *PhDsx* transcripts in both sexes.

Predicted proteins encoded by all three transcripts share the DM domain at their N-termini. However, we failed to detect an oligomerization domain in any of the *RpDsx* transcripts using either NCBI’s CDD/SPARCLE domain predictor software (67) or EMBL’s InterPro (71). We were similarly unable to detect an oligomerization domain in the Dsx protein sequences of another hemipteran – the planthopper *N. lugens* (58) – raising the possibility that hemipteran Dsx has lost this domain and thus has significant functional differences from the Dsx transcription factors of holometabolous insects (58).

#### Blattella germanica

With a combination of PacBio isosequencing and RACE, we recovered three *dsx* isoforms from *B. germanica*, each with a different terminal 3’ exon and 5’ and 3’ UTRs (Figure 2C). Like *dsx* genes described from the holometabolous orders Hymenoptera, Coleoptera, Lepidoptera, and Diptera (31,32,55,72), these isoforms are identical at their N-termini but differ by alternative splicing at the C-termini. The shared N-terminal sequence contains both the DM domain and the oligomerization domain (Figure 2C). RT-PCR and qPCR revealed that *BgDsx1* is found only in males, while *BgDsx2* and *BgDsx3* are female-specific (Figure 2D). This pattern of sexually dimorphic splicing, in which male and female transcripts share the DM domain but have different C-terminal domains, is typical of *dsx* in holometabolous insects (32,37,73).

#### Pediculus humanus

5’ and 3’ RACE identified two *PhDsx* isoforms, *PhDsx1* and *PhDsx2*, both of which were found in both male and female RACE libraries (Figure 2E). While both *PhDsx1* and *PhDsx2* have DM domains at their N-termini, alternative exon inclusion at the C-terminus yields a predicted Dsx dimerization domain in *PhDsx2* but not *PhDsx1*. RT-PCR on a different set of male and female louse samples confirmed that both *PhDsx1* and *PhDsx2* were expressed in both sexes (Figure 2F).

The order Blattodea, which includes cockroaches, is the most basal insect group in which *dsx* splicing has been examined to date. Thus, our results suggest that sex-specific *dsx* splicing appeared early in insect evolution, and the sharing of isoforms between sexes in P. humanus may represent a secondary loss of sex-specificity.

### *tra* is necessary for female, but not male development in *B. germanica*

*B. germanica* has a number of overt sexually dimorphic traits. Adult females are darker and have a rounded abdomen, while male abdomens are more slender (Figure 3A). Males also have a tergal gland that produces a mixture of oligosaccharides and lipids upon which females feed prior to copulation (74). This gland, located on the dorsal abdomen under the wings, appears externally as invagination in the 7^th^ and 8^th^ tergites, resulting in a complex structure with depressions and holes that lead to the internal glands (Figure 3B) (75). Finally, the ovaries, testes, and accessory glands that compose the male and female reproductive systems can be easily distinguished upon dissection (Figure 3C, D). To test whether *BgTra*, with its novel splicing pattern, functions to produce male and female traits similarly to its holometabolous orthologs, we used RNAi to knock it down at different stages of development.

In separate experiments, we injected female and male 4^th^, 5^th^ and 6^th^ instars with two overlapping double-stranded RNA sequences common to all *BgTra* isoforms (Figure 1B). *BgTra* expression levels were reduced to similar, very low levels in both males and females despite starting from a higher baseline level in females (Supplemental Figure 6). Females treated with dsBgTra developed normally until the 6^th^ instar, which took 16 – 21 days instead of the normal 8 days. Sixth instar female nymphs had an abnormal extrusion of tissue at the level of the last abdominal sternite (Supplemental Figure 7). Like wild-type females, dsBgTra females had five visible abdominal sternites, but the last sternite had an abnormal morphology with a “cut away” shape through which soft tissue was visible (Supplemental Figure 7). dsBgTra female nymphs molted into masculinized adults (Figure 3). All dsBgTra-treated female nymphs that molted to adults (n = 25) showed the slender abdomen tapered at the posterior end, which is typical of males (Figure 3A), and all had well developed tergal glands (Figure 3B). To test whether the tergal glands of masculinized females were functional, we extracted the oligosaccharides from the tergal glands of 11 dsBgTra females and 10 wild-type adult males and analyzed them by mass spectrometry. All of the 26 oligosaccharides present in the wild-type male tergal glands were also observed in dsBgTra female adults; these BgTra-depleted adults did not have any additional oligosaccharides not found in wild-type adult males (Supplemental Figure 8).

**Figure 3:**
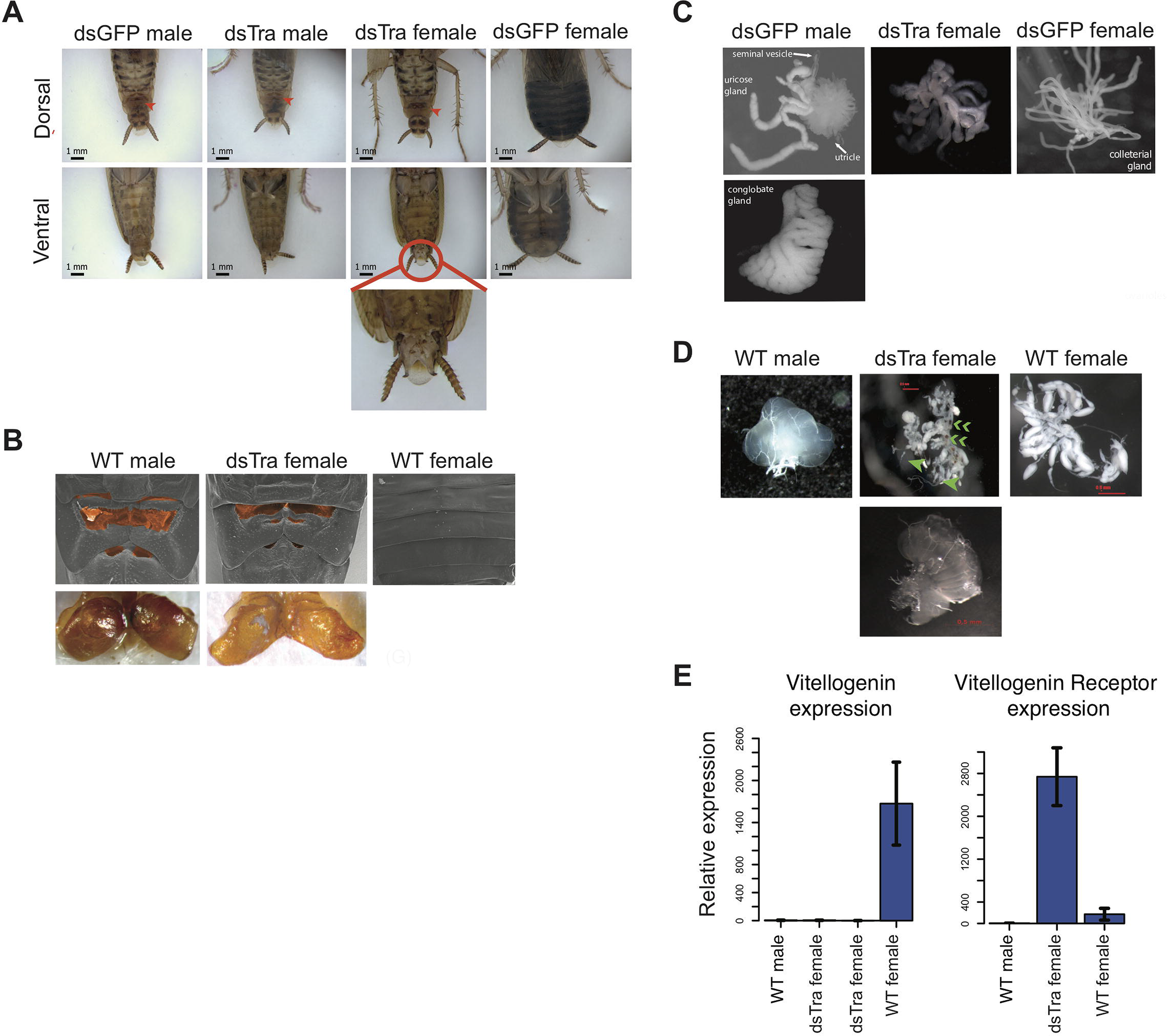
*BgTra* is necessary for female-specific but not male-specific sexual differentiation in *Blattella germanica*. (A) dsBgTra females, injected with *dsBgTra* in their 4th, 5th, and 6th instars, molt to adults with a masculinized abdomen, in contrast to control animals injected with dsGFP. Top row, dorsal view. Bottom row, ventral view. The tergal gland in wild-type males and dsBgTra females is indicated with red arrows. Red circle outlines abnormal tissue extrusion typical of dsBgTra females; zoomed-in view of this tissue is shown below. (B) Scanning electron microscopy (top row) and light microscopy (bottom row) reveal developed male-like tergal glands (shaded orange in SEMs and dissected for light microscopy) in dsBgTra females compared to wild-type (WT) females. (C) dsBgTra females have malformed, partly masculinized colleterial glands. Left column shows wild-type male accessory glands composed of seminal vesicles, utricles, uricose glands, and a conglobate gland. Right column, wild-type female colleterial gland. Center column, gland dissected from dsBgTra female is located in the same position as the wild-type colleterial gland, but shows thickened tubules reminiscent of the male accessory glands. (D) dsBgTra female gonads are deformed or intermediate between male and female gonads. Left column shows testis of a 5-day-old wild-type male. Middle column, top image shows disorganized gonad tissue in a typical 2-week-old dsBgTra female. Most individuals (n=12) had underdeveloped ovaries with some undifferentiated tissue, some ovarioles (green arrows), and some sclerotized cuticle (green chevrons) not normally found in the ovary. Middle column, bottom shows that some dsBgTra females (n=3) have intersex gonads composed of both ovarian and testicular tissue (indicated by green arrow). Right column shows ovaries of a 2-day-old wild-type female. (E) *vitellogenin*, an egg yolk protein, production is under the control of *BgTra*. Left: qPCR measuring relative vitellogenin expression in the fat body of males and females treated with dsBgTra. Right: relative upregulation of vitellogenin receptor transcription in the gonads of dsBgTra females, presumably due to the lack of circulating vitellogenin. *vitellogenin* and vitellogenin receptor gene expression was normalized to actin.

Internally, most adults that molted from the dsBgTra-treated female nymphs had underdeveloped ovaries containing large areas of undifferentiated tissue (Figure 3D). Some dsBgTra females had intersexual gonads (“ovotestes”) containing both ovarian and testicular tissue (Figure 3D). The colleterial gland, which is located at the base of the abdomen and is responsible for egg case protein production, was also partly masculinized, displaying thickened tubules reminiscent of male accessory glands (Figure 3C).

In contrast, male nymphs subjected to an equivalent *BgTra* RNAi treatment progressed normally through development. All dsBgTra male nymphs molted into adult males with wild-type external and internal morphology (n = 20); these adults were fertile, producing broods of normal size and number.

In *Drosophila*, the sex determination pathway regulates the production of sex-specific cuticular hydrocarbons (CHCs) (76,77). We find that in *B. germanica*, male and female CHC production is controlled by *BgTra*. The CHC profile of dsBgTra females is more similar to that of wild-type males than wild-type females; in particular, the precursor of the female contact sex pheromone is depleted in dsBgTra females, while the abundance of male-biased CHCs is increased (Supplemental Figure 9). dsBgTra females have significantly more total CHCs than either wild-type males or wild-type females (Supplemental Figure 9), a phenomenon likely explained by their non-functional ovaries. In wild-type females, large amounts of hydrocarbons are provisioned to the maturing oocytes (78). In ovariectomized wild-type females, some of the excess hydrocarbons which were destined to provision the ovaries are shunted to the cuticle (79); a similar process may be at work in dsBgTra females.

In addition to its female-specific effect on external and internal morphology, *BgTra* RNAi had a female-specific effect on gene expression. Expression of vitellogenin (yolk protein) in the fat body is high in wild-type adult females and absent in males. In dsBgTra females, vitellogenin expression was reduced to undetectable levels, similar to wild-type males (Figure 3E). Perhaps in response to the shortage of vitellogenin, expression of the vitellogenin receptor was upregulated in the gonads of dsBgTra females (Figure 3E). Overall, we conclude that *BgTra*, like its holometabolous insect orthologs, is required for female sexual differentiation but is dispensable in males.

### *tra* represses male-specific behavior in female cockroaches

In *D. melanogaster*, sex-specific behavior is indirectly under the control of *tra* via the *fruitless* (*fru*) transcription factor (80–82). *fru* orthologs also control courtship behavior in other Dipterans and in *B. germanica*, and *fru* shows conserved sex-specific splicing across holometabolous insects (83–85). In *D. melanogaster* females, Tra prevents expression of *fru* isoforms that direct male courtship behavior (81). To test whether *BgTra* also controls sex-specific behavior in *B. germanica*, we exposed ten masculinized dsBgTra female adults to antennae clipped from sexually mature wild-type females. In typical courtship, wild-type males are stimulated by a contact sex pheromone on the female antenna. Upon contact with a female antenna, wild-type males orient their abdomen towards the antenna and raise their wings to display the tergal glands. As the female feeds on tergal gland secretions, the male extends his abdomen, grasps the female’s genitalia, and copulation follows (86). Wild-type females do not show the wing-raising behavior. Nine out of ten masculinized dsBgTra female adults showed the same response as wild-type males, although the mean lag time between introduction of the female antenna and wing raising was longer in dsBgTra females (mean = 19.9 seconds) than in wild-type males (mean = 4.9 seconds) (Welch’s t-test, p = 0.02) (Figure 4 A, B). These results indicate that dsBgTra females, unlike wild-type females, perceive wild-type female sex pheromone and respond with courtship behavior. Similar to its function in holometabolous insects, *BgTra* is required to repress male-specific behavior in *B. germanica*.

**Figure 4:**
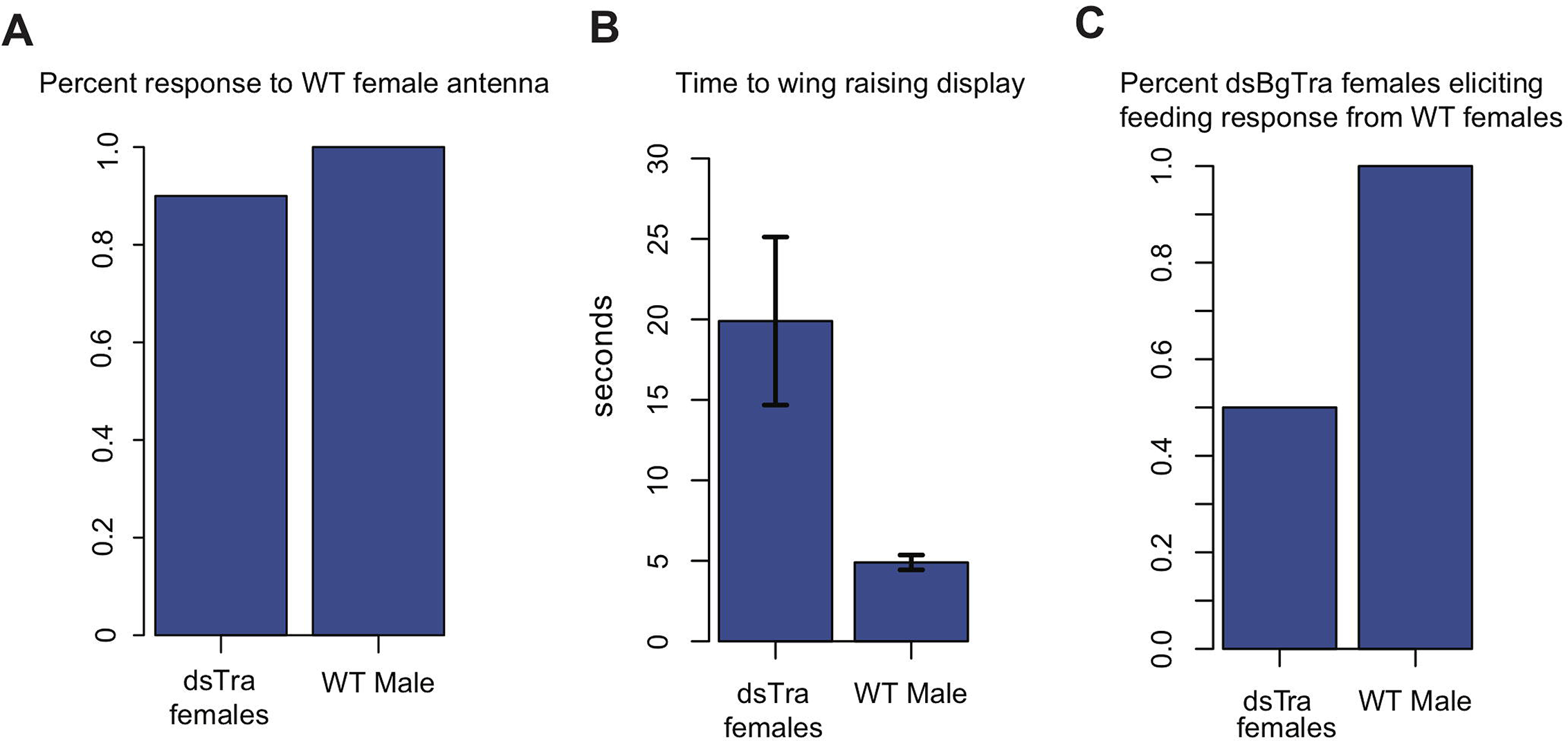
dsBgTra females perform male courtship behavior and elicit courtship responses from wild-type females. In courtship, wild type male *Blattella germanica* raise their wings to display their tergal gland. When courtship is successful, wild type females feed upon the secretions of this gland prior to copulation. dsBgTra females (n=10) and wild-type males (n=10) were exposed to an antenna clipped from a wild-type, 7 day old virgin female, and their response times (in seconds) were compared. (A) Nine out of ten dsBgTra females performed the stereotypical male wing-raising courtship display in response to a female antenna compared to 0/10 of wild-type females. (B) However, dsBgTra females took longer than wild-type males to respond with the wing-raising display. (C) 5/10 dsBgTra females elicited feeding response from wild-type females after raising their wings.

In a separate set of experiments, we tested whether wild-type females respond to the wing-raising display of masculinized dsBgTra females. In five out of ten instances, wild-type females began feeding on the tergal glands of dsBgTra females, confirming the functionality of these glands (Figure 4 C). Overall, the results indicate that dsBgTra females express male sexual traits that attract wild-type females.

### *BgTra* is necessary for female, but not male embryonic viability

We also found that *BgTra* is crucial for female embryonic development. Three-day-old adult females injected with dsBgTra produced all-male broods (Figure 5A), suggesting that maternal dsBgTra treatment either masculinizes or kills female progeny. Female-specific lethality appears more likely for two reasons. First, the number of offspring in the all-male broods from dsBgTra females was about 50% of the offspring from dsGFP-injected control females (Figure 5B). Second, all the broods that hatched from dsBgTra mothers contained dead embryos remaining in the ootheca (Figure 5C). One possibility is that *BgTra* may affect dosage compensation in the cockroach, which has male heterogametic, XX/XO sex determination (87). When *T. castaneum* females are injected with dsRNA targeting *tra*, they also produce all-male broods with much reduced survival (36). These results suggest that *tra* could either have a deeply conserved role in dosage compensation, or has repeatedly evolved a role in this process.

**Figure 5:**
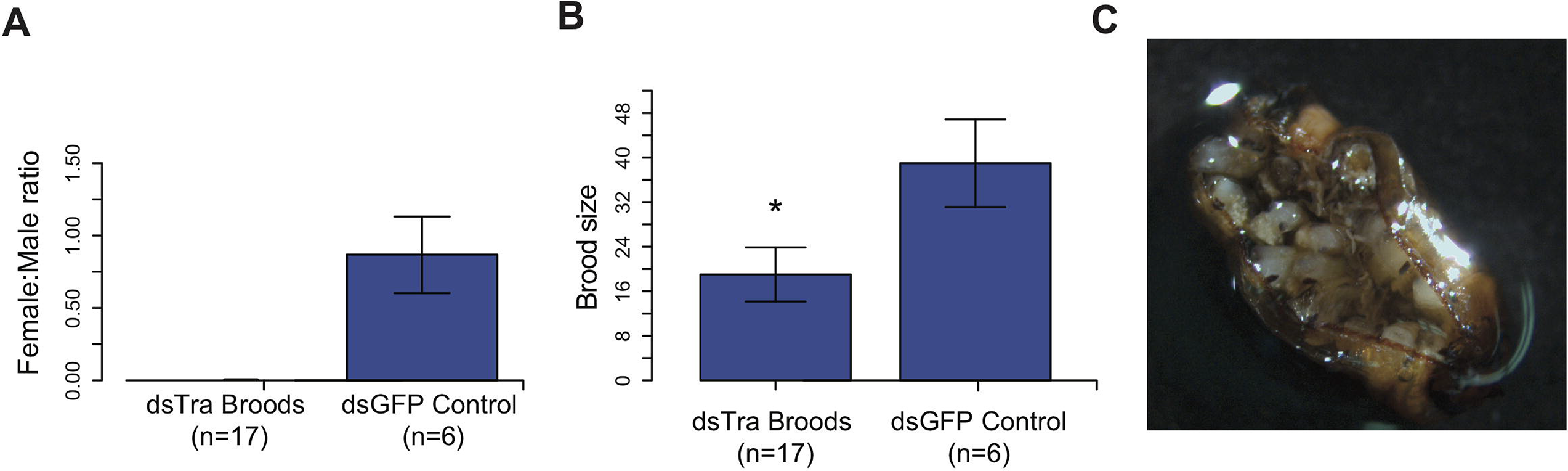
Maternal knockdown of *BgTra* in *Blattella germanica* results in all-male broods.. (A) produced all-male broods, compared to control females injected with dsBgGFP (n=6) (B) Brood sizes from mothers injected with dsBgTra were significantly smaller than the dsGFP controls (Welch’s t-test, p=0.001). (C) Image of a typical ootheca (egg case) from a dsBgTra mother that failed to hatch. Of 22 mothers injected with dsTra, 5 egg cases failed to hatch completely. The remaining oothecae had 5 – 19 dead embryos remaining within it after all males had emerged.Variability in the number of dead embryos can be attributed to surviving offspring eating dead embryos, which was observed by the authors.

### *B. germanica BgTra* controls sex-specific splicing of *BgDsx*

In *B. germanica*, *BgDsx* has two female-specific and one male-specific isoform (Figure 2C). Because *dsx* splicing is under the control of *tra* in most holometabolous insects, we tested whether *BgTra* RNAi affected the production of sex-specific *BgDsx* isoforms. qPCR and RT-PCR revealed that in dsBgTra females, *BgDsx* splicing switched completely from the female to the male pattern (Figure 6). Neither of the two female-specific transcripts were present in dsBgTra females, whereas the male-specific *BgDsx* isoform was present in abundance similar to wild-type males (Figure 6). This is identical to the pattern seen in holometabolous insects such as *D. melanogaster* and *T. castaneum* (36,65).

**Figure 6:**
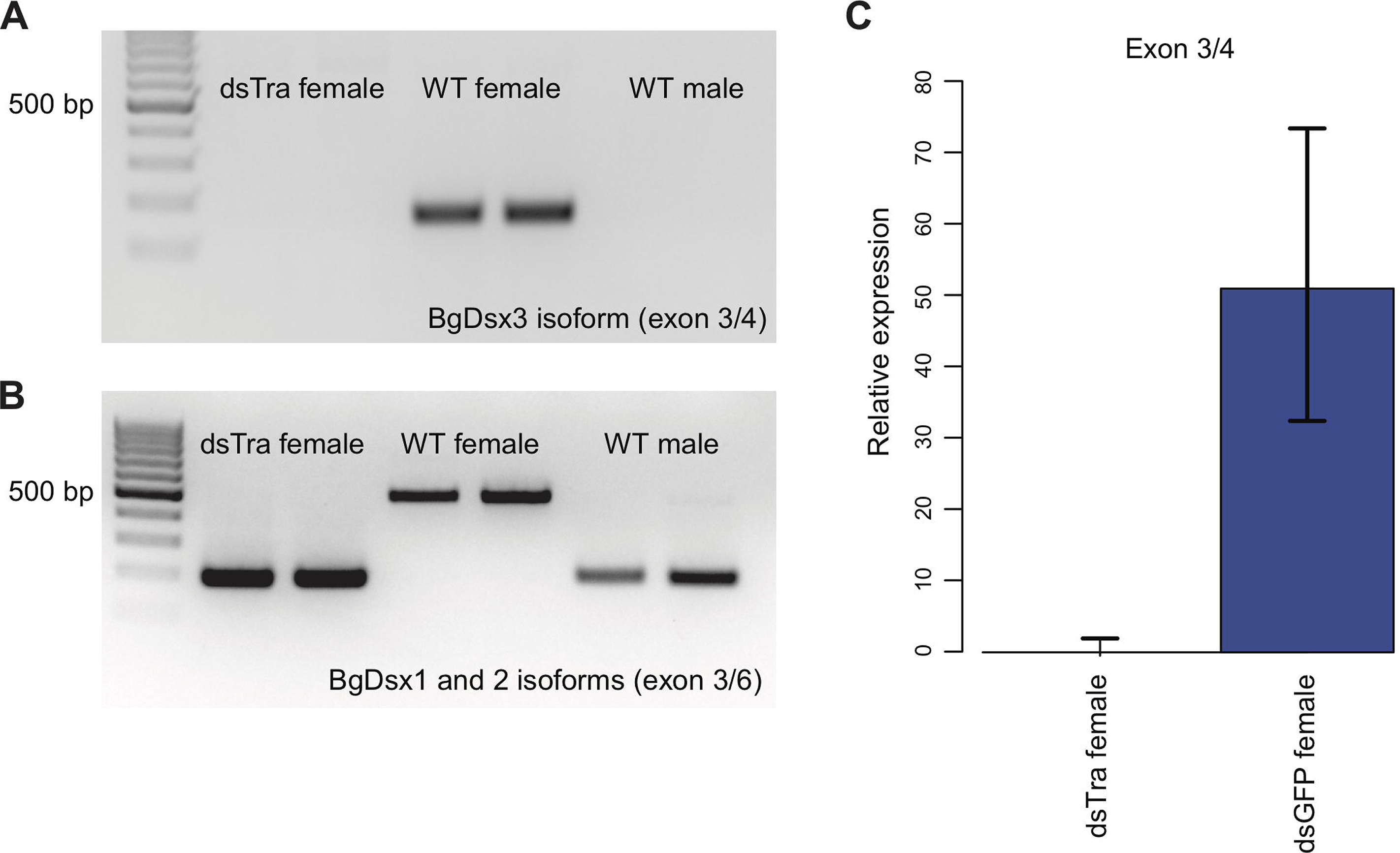
*BgTra* controls sex-specific splicing of *BgDsx*. (A) RT-PCR showing that the female-specific *BgDsx* exon 3/4 junction (Fig. 2 B,C) is absent in dsBgTra females (B) RT-PCR showing that the male-specific *BgDsx* exon 3/5 junction (Fig. 2B) is present in dsBgTra females. dsBgTra females do not express any of the female-specific *BgDsx* isoforms. (C) qPCR shows complete absence of the female-specific *BgDsx* exon 3/4 junction in dsBgTra females.

### *BgDsx* is necessary for male-specific but not female-specific sexual differentiation

In separate experiments, we injected three different dsRNAs targeting *BgDsx*. Two of these dsRNAs targeted the DNA-binding domain shared by male and female *BgDsx* transcripts; a third dsRNA targeted the Dsx Dimerization Domain, also shared between males and females (Figure 2). Surprisingly, we observed that *BgDsx* transcript abundance increased after injection with dsBgDsx in both males and females (Supplemental Figure 10), making it difficult to measure the efficiency of RNAi treatment. Increased expression of a target gene after dsRNA treatment has been previously described (88), and there is not always a strong relationship between transcript abundance and RNAi phenotype (89). Most plausibly, the upregulation observed after a dsRNA treatment results from a rebound effect on the transcription of the targeted gene. Although the effect of RNAi on *BgDsx* transcript abundance was similar in males and females, its effect on adult morphology and downstream gene expression was markedly different between the two sexes.

Both males and females injected with dsBgDsx during 4^th^, 5^th^, and 6^th^ instars progressed normally through nymphal development. However, upon the adult molt, all males (n = 23) had an abnormal extrusion of tissue at the tip of their abdomen (Figure 7A). This extrusion appears to be an outgrowth of the ejaculatory duct (Supplemental Figure 11). A range of body shapes was observed in these adults, from slender abdomens typical of wild-type males to the rounder abdomens typical of females. The tergal glands of dsBgDsx males also ranged from fully developed to severely reduced (Figure 7A). Moreover, the dsBgDsx males showed darker pigmentation, similar to wild-type females, especially on the dorsal side (Figure 7A). Because of their malformed ejaculatory ducts, dsBgDsx males were not able to mate, making it impossible to assess their fertility. Dissection of dsBgDsx males revealed apparently normal testes (n = 15), with morphologically normal sperm indistinguishable from wild-type *B. germanica* sperm (n = 3). However, the conglobate glands of dsBgDsx males failed to mature over the first week of adult life, while tissue discoloration was observed in the utricles (Supplemental Figure 12).

**Figure 7:**
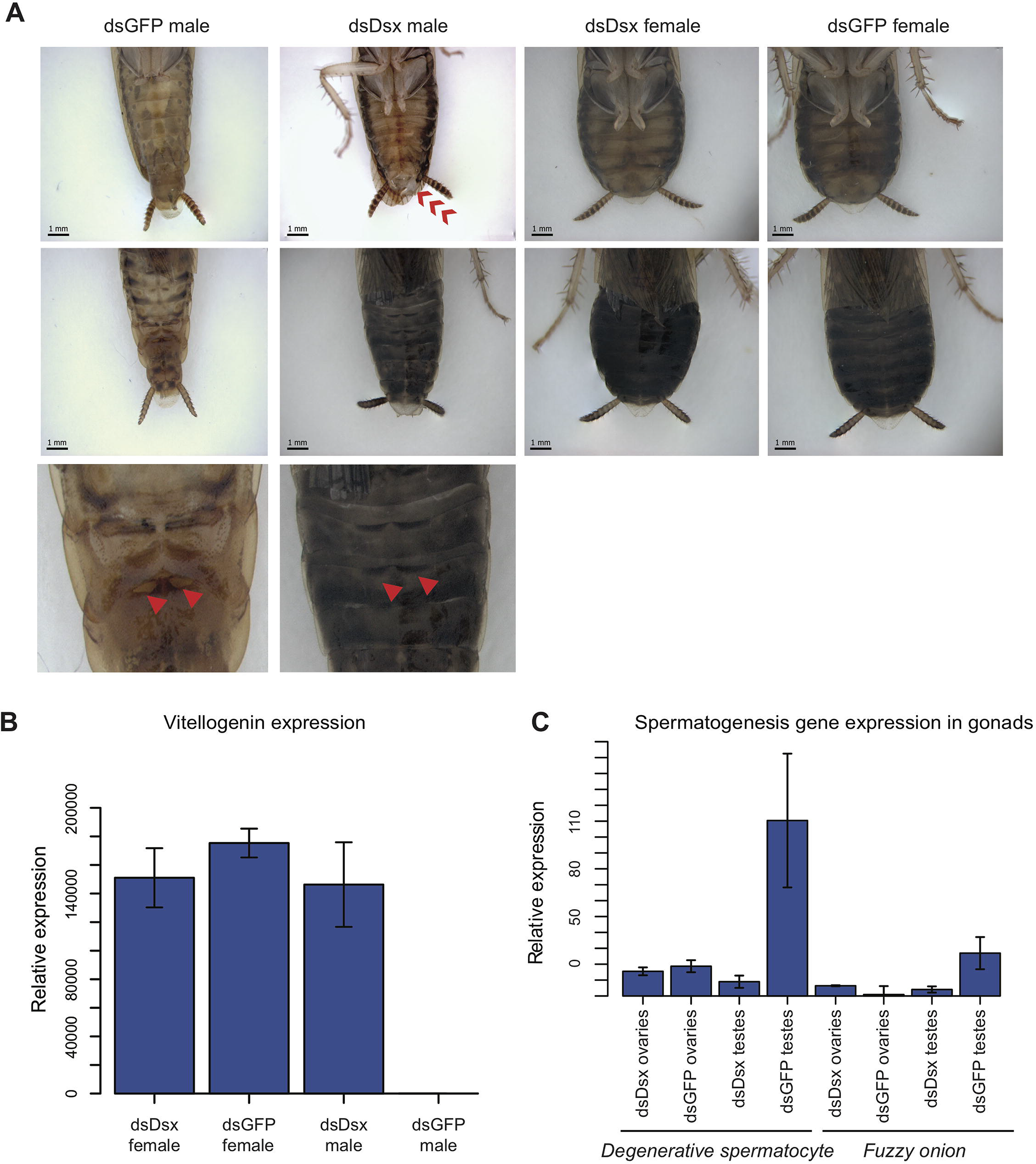
dsBgDsx RNAi in *Blattella germanica* feminizes males but has no effect in females. (A) dsBgDsx males have reduced tergal glands (red arrow), darker female-like pigmentation, and an extrusion of tissue at the distal end of the abdomen (red chevron) compared to dsGFP injected males (see also Supplemental Figure 12). Females are unaffected by dsBgDsx treatment. Bottom row shows zoomed-in view of dorsal segments containing the tergal gland. The openings from which tergal glands secrete their oligosaccharides and lipids in dsGFP males (red arrows) are greatly reduced in dsBgDsx males. (B) BgDsx represses *vitellogenin* expression in males, but has no effect in females (n=3 biological replicates per treatment). (C) BgDsx controls the expression of two spermatogenesis-related genes in males, but has no effect on their expression in females. qPCR showing the expression of two genes important for *Drosophila melanogaster* spermatogenesis, *degenerative spermatocyte* and *fuzzy onions*, relative to *actin* (n=3 biological replicates for all columns).

dsBgDsx females molted into adults with no visible external abnormalities (Figure 7A), and had apparently normal gonads and reproductive tracts. Adult dsBgDsx females elicited courtship from wild-type males, mated with them, and produced broods whose size was not significantly different from those of control females (Supplemental Figure 13). When virgin females received dsBgDsx injections 3 days before mating, they produced broods of normal size with normal sex ratio (Supplemental Figure 13). Thus, it appears that BgDsx controls many though not all male-specific traits, but has no obvious effect on female-specific development. This is clearly different from holometabolous insects, where *dsx* plays active roles in both male and female sexual differentiation, and where *dsx* mutants develop into intersexes that are intermediate in phenotype between males and females (51).

To test whether BgDsx controlled sex-specific gene expression, we examined the expression of two genes, *fuzzy onions* (*fzo*) and *degenerative spermatocyte* (*des*), that function in spermatogenesis in *D. melanogaster* and have homologs throughout metazoans (90–93). Expression of both genes was greatly reduced in the testes of dsBgDsx males compared to the testes of wild-type males, whereas no significant effect was seen in the ovaries of dsBgDsx females (Figure 7C). Conversely, the female-specific *vitellogenin* gene was strongly upregulated in the fat body of dsBgDsx males compared to wild-type males, reaching the level normally seen in wild-type females (Figure 7B). In contrast, vitellogenin expression was not affected in dsBgDsx females (Figure 7B). Together, these results suggest that *BgDsx* controls downstream gene expression in males but not in females. This pattern, while different from holometabolous insects, has been observed in the hemipteran *N. lugens*, where a *dsx* ortholog has a female-specific isoform that appears to play no role in vitellogenin production (58). In *D. melanogaster*, yolk protein genes are upregulated by the female-specific isoform of *dsx* and repressed by the male-specific isoform, so that *dsx* mutants show an intermediate level of yolk protein gene expression (94).

## DISCUSSION

The sex determination pathway must by necessity produce a bimodal output: male or female. How this output is achieved varies considerably across animal taxa and can rely on molecular processes as different as systemic nuclear hormone signaling, post-translational protein modification, or, in the case of insects, sex-specific alternative splicing. In this paper, we examined sexual differentiation in hemimetabolous insects to investigate the evolutionary origin of this unique mode of sexual differentiation.

The canonical *transformer*-*doublesex* pathway of holometabolous insects has three derived features that are not found in non-insect arthropods or in non-arthropod animals: (1) the presence of distinct male- and female-specific splicing isoforms of *dsx* that play active roles in both male and female sexual development; (2) the production of functional Tra protein in females but not males; and (3) the control of female-specific *dsx* splicing by Tra. Our work in hemimetabolous insects suggests that these elements evolved separately and at different times, and that the definitive *tra*-*dsx* axis was assembled in a stepwise or mosaic fashion.

### Sex-specific splicing of *doublesex* predates its role in female sexual differentiation

Distinct male and female *dsx* isoforms have been reported in all holometabolous insects examined, as well as in the planthopper *N. lugens* (Hemiptera) (58). Our results confirm the presence of male and female *dsx* isoforms in another hemipteran, *R. prolixus,* and show that this pattern is conserved through Blattodea – the most basal insect order studied to date. This suggests that sex-specific splicing of *dsx* was likely the first feature of the canonical insect sexual differentiation pathway to evolve. In *P. humanus* (Phthiraptera), the alternative splicing of *dsx* follows the usual pattern, with a common N-terminal domain and alternative C-termini, but we found that both *dsx* isoforms are present in both sexes. Previous work in the hemipteran *Bemisia tabaci* also failed to detect sex-specific *dsx* isoforms (95), suggesting that some insects may have secondarily lost sex-specific *dsx* splicing.

We did not detect any role for the cockroach *dsx* gene in female sexual differentiation in *B. germanica*. Recent work in the Hemipteran *N. lugens* produced similar results: despite the presence of male and female *dsx* isoforms, *N. lugens* females developed normally following *dsx* RNAi knockdown, while the males were strongly feminized (58). Together, these results suggest that in at least some hemimetabolous insects, *dsx* is spliced as in holometabolous insects, but may function as in the crustacean *Daphnia*, where *dsx* is necessary for male but not female development (53).

The existence of a female-specific *dsx* isoform in the absence of a female-specific *dsx* function is, of course, surprising. Although we examined multiple features of female external and internal anatomy, gene expression, behavior, and reproduction in *B. germanica* dsBgDsx females, we cannot rule out that *dsx* performs some subtle or spatially restricted function in female cockroaches. Such relatively minor and tissue-specific role, if it exists, could provide a crucial stepping stone in the origin of the classical *dsx* function as a bifunctional switch that is indispensable for female as well as male development. One possibility is that, similar to other transcription factors, *dsx* originally evolved alternative splicing as a way of promoting the development of different cell types. If some *dsx* isoforms evolved functions that were dispensable for male sexual development, *dsx* could gradually lose its ancestral pattern of male-limited transcription, opening the way for the evolution of new isoforms with first subtle, but eventually essential, functions in females.

### The role of *transformer* in sexual differentiation predates canonical sex-specific splicing of tra

The opposite pattern is observed for *transformer,* which has apparently evolved a sex-specific role before that role came to be mediated by sex-specific splicing. In most holometabolous insects, alternative splicing of *tra* produces a functional protein in females but not in males, due to a premature stop codon in a male-specific exon near the 5’ end of *tra*. In hemimetabolous insects, we only observe this type of sex-specific splicing in *R. prolixus*. In *B. germanica* and *P. humanus*, full-length and presumably functional Tra proteins that include all of the important functional domains are produced in both males and females. Despite this, we find that the cockroach *tra* gene has a female-limited function in sexual differentiation, and appears to be completely dispensable for male development – just as it is in holometabolous insects. Thus, the function of *tra* in female sexual differentiation predates the evolution of female-limited Tra protein expression.

Although they lack the canonical sex-specific splicing observed in holometabolous insects, the *tra* genes of *B. germanica* and *P. humanus* show a different pattern of alternative splicing affecting the CAM domain, which plays a role in *tra* autoregulation of holometabolous insects (62). In some *tra* isoforms, read-through of an exon containing the 5’ half of the CAM domain results in a premature stop codon; interestingly, these truncated isoforms are more abundant in female *B. germanica* than in males; in *P. humanus*, these transcripts are found in both sexes. In *BgTra*, isoforms with an intact reading frame come in two types. Half of these intact isoforms contain an in-frame exon spliced into the middle of the CAM domain; in the others, this exon is spliced several base pairs upstream of the stop codon, producing an uninterrupted CAM domain. This particular splice junction in the middle of the CAM domain is conserved in holometabolous insects (61), and the amino acid residues spanning this junction are necessary for female-specific autoregulation of *tra* in the housefly (62). In *P. humanus*, we only isolated *tra* isoforms with an exon interrupting the CAM domain. It appears that the evolution of the canonical holometabolous splicing of *tra*, with a male-specific premature stop codon, coincided with a loss of *tra* isoforms with an interrupted CAM domain. In fact, the splicing pattern of *tra* in *R. prolixus* suggests that the male-specific stop codon has originally evolved in the middle of the CAM domain (Figure 1A).

### The role of *tra* in regulating *dsx* splicing predates sex-specific expression of Tra protein

Although the function of *tra* in female sexual development in *B. germanica* does not appear to be mediated by *dsx*, we find that *tra* is necessary for sex-specific *dsx* splicing in the cockroach, as it is in all holometabolous insects except Lepidoptera (96). The mechanism of *dsx* regulation by *tra* is likely to be different in *B. germanica* compared to holometabolous insects. In the latter, functional Tra protein is absent in males but present in females, where it dimerizes with the RNA-binding protein Tra-2 to regulate the splicing of *dsx* and *fru* (68,68,97). In the cockroach, however, full-length Tra proteins containing all the functional domains are expressed at similar levels in both sexes, so that male-specific splicing of *dsx* in males cannot be explained by lack of Tra. It could be due instead to an interaction of Tra with different binding partners in males vs females. We note that while the *D. melanogaster* Tra protein does not contain an RNA-binding domain and does not bind to RNA without Tra-2, the cockroach Tra protein, as well as those of some other insects and crustaceans, contain predicted RNA recognition motifs (65). Thus, it is possible that Tra first evolved its function in regulating alternative splicing as a broadly acting RNA-binding protein that worked in concert with other splicing factors, before losing its ability to bind RNA and becoming a dedicated partner of Tra-2 with a narrow range of downstream targets. If so, the situation we find in *B. germanica* may represent a transitional stage in the evolution of the canonical *tra*-*dsx* axis, where *dsx* is one of many Tra targets, rather than the main mediator of its female-specific function. In this scenario, the *tra*-*dsx* axis evolved via merger between expanding *dsx* function (from males to both sexes) and narrowing *tra* function (from a general splicing factor to the dedicated regulator of *dsx*).

### Evolution of the insect sexual differentiation pathway: stepwise or mosaic?

We propose that the canonical insect mechanism of sexual differentiation based on sex-specific splicing of *dsx* and *tra* evolved gradually in hemimetabolous insects, and may not have been fully assembled until the last common ancestor of the Holometabola (Figure 8). In crustaceans, mites, and non-arthropod animals, *dsx* and its homologs act as male-determining genes that are transcribed in a male-specific fashion and are dispensable for female sexual differentiation (53,54,57). One of the earliest events in insects was the evolution of sex-specific *dsx* splicing, where *dsx* is transcribed in both sexes but produces alternative isoforms in males vs females. We do not know whether Tra was involved in sex-specific *dsx* splicing from the beginning, or evolved this function later; in either case, the function of *tra* in controlling female sexual differentiation may well predate its role in regulating *dsx* splicing. Eventually, however, Tra became essential for the expression of female-specific *dsx* isoforms. At the next step, expression of functional Tra protein became restricted to females due to the origin of a male-specific exon with a premature stop codon at the 5’ end of the *tra* gene. As this process unfolded, female-specific *dsx* isoforms gradually evolved female-specific functions, which may have been minor at first but became essential at or before the origin of holometabolous insects (Figure 8).

**Figure 8:**
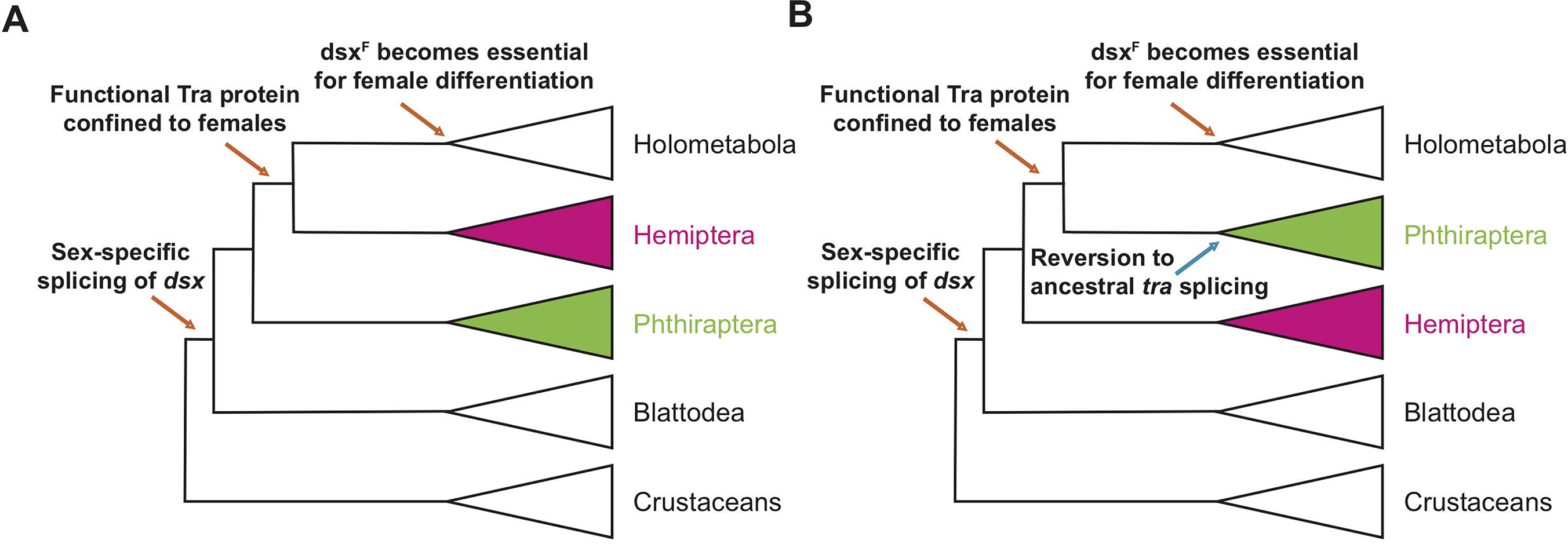
Evolutionary assembly of the insect sexual differentiation pathway. Uncertainty in therelationships among Hemiptera, Phthiraptera, and Holometabola (Li et al, 2015; Freitas et al, 2018; Misof et al, 2014, Johnson, 2018) means that the sequence of events in the evolution of the *tra-dsx* axis is unclear. If Hemiptera (pink) are the sister group of Holometabola, the key molecular innovations have evolved in a stepwise order (A). If Phthiraptera (green) are the sister of Holometabola instead, the evolution of the *tra-dsx* axis must have followed a more mosaic pattern, with at least some secondary reversions (B).

The details of this model depend on the phylogenetic relationships of hemimetabolous insect orders and the holometabolous clade, which remain controversial. Despite rapidly growing amounts of data, phylogenetic analyses have had limited success in identifying the closest outgroup to Holometabola. In some phylogenies, that outgroup is Hemiptera (Figure 8A) (98); in others it is Psocodea, which includes lice (Figure 8B) (99–101); in all molecular phylogenies to date, the relevant nodes are not strongly supported. In fact, some phylogenies show Hemiptera and Psocodea as a polytomy (102,103). The holometabolous splicing pattern of *tra* is seen in *R. prolixus* but not in the cockroach or louse, suggesting that either Hemiptera are the closest outgroup to holometabolous insects, or the splicing of *tra* in *P. humanus* represents a reversion to an ancestral state (Figure 8).

Our study of *transformer* and *doublesex* expression and function across hemimetabolous insects establishes a broad outline for the origin of the unique insect-specific mode of sexual differentiation via alternative splicing. Comparative and functional work in other basal insects, non-insect Hexapods, and crustaceans will be needed to determine the ancestral functions of *tra*, the roles of female-specific *dsx* isoforms in hemimetabolous insects, and other details of this emerging model.

## MATERIALS AND METHODS

### DMRT and SR family gene trees

To differentiate *dsx* orthologs from other DMRT paralogs, we made phylogenetic trees of arthropod DMRT genes. DMRT protein sequences from previously studied arthropods (Wexler and Plachetzki, 2014) (*D. melanogaster, A. mellifera, B. mori, T. castaneum, R. prolixus, D. magna,* and *I.s scapularis*) were used as queries (Supplemental File 1) in BLASTP searches of predicted gene models from 22 other arthropod species (Supplemental Table 1). Hits with e-values lower than 0.01 were retained and added to the file that included the original DMRT queries. This sequence file was multiply aligned with the MAFFT program using the –geneafpair setting (104). Upon alignment, sequences without the characteristic DMRT DNA-binding domain CCHHCC motif (104)were discarded, and the entire multiple alignment was rerun.

We used a similar procedure to identify *tra* orthologs in hemimetabolous insects. For the RS family gene tree, the query file consisted of Tra, Transformer-2, Pinin, and SFRS protein sequences from *D. melanogaster, B. germanica*, and *D. magna* (Supplemental File 2), and a BLASTP search was performed on the same arthropod species, retaining hits with e-values <0.01. After sequences were multiply aligned with the MAFFT algorithm, gaps in the alignment were trimmed with trimAl using the –automated1 setting (105).

To construct gene trees for DMRT and RS families, prottest3.4 (106) was used for model selection for the amino acid alignment; RAxML8.2.9 was used to build a maximum likelihood tree with a WAG+G model for both the SR family gene tree and the DMRT gene tree (107).

### Cockroach husbandry

Functional experiments were conducted with two German cockroach *B. germanica* strains: Orlando Normal, obtained from Coby Schal’s laboratory at North Carolina State University; and Barcelona strain from Xavier Belles’s laboratory at the Institute of Evolutionary Biology. *B. germanica* were kept in an incubator at 29°C and fed Iams dog food *ad libitum*.

### PacBio isosequencing

Pacific Biosystems (PacBio) isosequencing was performed separately on male and female *B. germanica* tissues. We extracted RNA from reproductive organs (gonad and colleterial glands) and fat body of a seven day old virgin female and from reproductive organs (gonad, conglobate gland, accessory glands, and seminal vesicles) and fat body of two seven day old males for library preparation. Tissues were pooled in Trizol, and a separate RNA extraction was done for each sex. Clontech SMARTER PCR cDNA synthesis kit and PrimeSTAR GXL DNA polymerase were used for cDNA synthesis; a cDNA SMRTbell kit was then used to add sequencing adapters. The male and female libraries were run on separate SMRT cells on the PacBio Sequel instrument, and the PacBio SMRTLink software was used for downstream data processing. We used tBLASTn to search the high and low quality polished consensus isoforms for *BgTra* and *BgDsx* transcripts. We obtained additional isoforms searching the sets of circular consensus sequences. All splice junctions from *BgTra* and *BgDsx* Isosequencing-generated transcripts were verified with RT-PCR. When the coding sequence of PacBio-generated isoforms disagreed with the *B. germanica* genome, sequence was verified by Sanger sequencing.

### Illumina RNA sequencing

To identify *dsx* and *tra* isoforms in *R. prolixus* and *P. humanus*, we used Trizol to extract RNA according to the manufacturer’s instructions. For *R. prolixus*, we extracted RNA separately from male and female gonad tissue. For *P. humanus*, adults were sexed and pooled before RNA extraction. For both insects, 1 µg of RNA was used to construct each library using the NEBNext kit and Oligos. Libraries were quantified with NEBNext Quant Kit and sequenced using 150 bp paired-end reads on an Illumina HiSeq4000 machine. Poor quality reads were discarded with FastQC (default parameters) (https://www.bioinformatics.babraham.ac.uk/projects/fastqc/) and trimmed with trimmomatic 0.36 using suggested filters (LEADING:3 TRAILING:3 SLIDINGWINDOW:4:15 MINLEN:36) (108). Trinity 2.1.1 with default parameters was used for *de novo* assembly of the male and female transcriptomes (109).

## 5’/3’ RACE

The Clontech 5’/3’ SMARTer Rapid Amplification of cDNA Ends (RACE) kit was used to make RACE libraries from male and female *R. prolixus*, *P. humanus*, and *B. germanica* total RNA. All RNA extractions were performed using Invitrogen TRIzol Reagent according to the provided protocol. For *R. prolixus* and *B. germanica*, we combined gonad tissue, reproductive tract tissue, and fat body from a single male or a single female to make the male and female RACE libraries, respectively. *P. humanus* RNA was extracted from pools of between 6-10 whole adult males or females.

For all species, 5’ RACE cDNA fragments were amplified using gene-specific primers (Supplemental Table 5) and kit-provided UPM primers and Clontech Advantage 2 Polymerase. Thermocycling conditions were: 95℃ for 2 mins; 36 cycles of (95℃ for 30 sec, a primer-specific annealing temperature for 30 seconds, 68℃ for 2.5 minutes); 95℃ for 7 minutes. 3’ RACE cDNA fragments were amplified using single or nested PCRs with gene-specific primers and either Clontech Advantage 2 Polymerase or SeqAmp DNA Polymerase (Supplemental Table 5). Clontech Advantage 2 PCRs were performed using the above cycling conditions, and SeqAmp DNA Polymerase touchdown thermocycling conditions were: 5 cycles of (94°C for 30 sec, 72°C for 1-2 min); 5 cycles of (94°C for 30 sec, 68°C for 1-2 min); 35 cycles of (94°C for 30 sec, 66°C for 30 sec, 72°C for 1-2 min). Nested PCRs used diluted (1:50) outer PCRs as template, a nested gene-specific primer and the UPM short primer, and 25 cycles of (94°C for 30 sec, 65°C for 30 sec, 72°C for 2 min). All PCR products were gel-purified using Qiagen QIAquick or Macherey-Nagel NucleoSpin gel and PCR clean up kits. PCR products produced by Advantage 2 Polymerase were cloned into the Invitrogen TOPO PCRII vector and SeqAmp DNA Polymerase PCR products were cloned into linearized pRACE vector (provided with the SMARTer RACE kit) with an In-Fusion HD cloning kit. Between 6-10 colonies were Sanger sequenced per PCR reaction with M13F-20 and M13R sequencing primers.

### Reverse-transcription PCR tests of isoform sex-specificity

With *R. prolixus* and *P. humanus*, we found that cDNA synthesized with the Clontech SMARTer 5’/3’ RACE kit amplified well in reverse-transcription PCRs (RT-PCR). We synthesized 3’ RACE-ready cDNA from two different female and two different male total RNA samples that were not used for 5’/3’ RACE. To test isoforms for sex-limited expression, we amplified isoform-specific sequences in female and male samples to using 40 cycles of PCR and SeqAmp DNA Polymerase. To test for sex-specificity of isoforms in *B. germanica*, we isolated total RNA from reproductive tracts and fat bodies of male and female adult roaches using Invitrogen TRIzol reagent according the manufacturer’s instructions. We then treated these RNA samples with Promega RQ1 DNaseI, also according to manufacturer’s instructions, and primed the RNA for cDNA synthesis using a 1:1 mix of oligo dT:random hexamers. We used Invitrogen SuperScriptIII for reverse transcription, and AccuPower Taq from Bioneer for PCR.

### Reverse-transcription qPCR

RNA for qPCR was isolated with Invitrogen TRIzol reagent and then treated with Promega RQ1 DNaseI in accordance with the manufacturer’s instructions for both reagents. Reverse transcription for qPCR was performed with Invitrogen SuperScriptIII or SuperScriptIV after RNA was primed with a 1:1 mix of oligo dTs and random hexamers. qPCR was done on non-diluted or diluted cDNA (1/5x) with BioRad Sso Advanced Universal SYBR Mix and on a BioRad CFX96 machine. We quantified expression in 2-3 samples per sex across three technical replicates each (we tested 3 females and 3 males from *B. germanica*, and 2 females 2 males from both *R. prolixus* and *P. humanus*). Target gene expression was normalized to β-*actin* expression using the Δ Ct method (110). A dilution series was used to test primer pairs for efficiency; only primer pairs with calculated efficiency between 90% - 110% were used.

### Protein organization

BgDsx domains were identified by NCBI domain predictor software: https://www.ncbi.nlm.nih.gov/Structure/cdd/wrpsb.cgi. For BgTra, the CAM domain was identified by visual inspection after a MAFFT alignment of BgTra protein sequences with previously studied Tra proteins in holometabolous insects (111). The RS domain was identified by calculating percent of protein sequence containing arginine and serine residues; the RRM domain was identified by NCBI domain predictor software.

### RNAi cloning and synthesis

Template for dsRNA targeting *BgTra* and *BgDsx* were amplified out of *B. germanica* cDNA using Bioneer AccuPower Taq. The following cycling parameters were used: 95°C for 2 minutes, then 35 x (95°C for 30 seconds, annealing temperature for 30 seconds, 72°C for 30 seconds), then a final extension of 72°C for 3 minutes. Templates were cloned into TOPO-TA PCRII from Invitrogen. Template for dsGFP was cloned with the same Taq, using the following cycling parameters: 95°C for 1 minute, then 35 x (95°C for 30 second, 55°C for 30 seconds, 72 for 30 seconds) following by an extension at 72°C for 3 minutes. Directional colony screens were performed to identify forward and reverse inserts which were combined, so that a single transcription reaction contained both forward and reverse templates. A Thermo Fisher Scientific MEGAscript RNAi kit was used to prepare dsRNA.

### RNAi injections, dissections and imaging

We conducted two sets of experiments for each gene in *B. germanica*. First, nymphs were injected throughout their development to assess the function of *BgTra* and *BgDsx* from the mid-point of juvenile development onwards. Second, virgin adult females were injected before mating in a parental RNAi experiment that tested for possible maternal roles of these genes during embryogenesis. For the first set of experiments, we sexed 4^th^ instar *B. germanica* nymphs and injected each individual once in the 4^th^ instar, once in the 5^th^ instar, and once in the 6^th^ instar. Fourth instar nymphs were injected with ∼500 ng of dsRNA in 0.5 µL; 5^th^ and 6^th^ instar nymphs females with 1 µg of dsRNA in 1 µL. Both nymphs and adults were injected between two sternites. For parental RNAi, we put injected females in individual containers with 2 – 5 wild-type adult males per container. Adult males were sacrificed after observing that mating had occurred (either by direct observation of mating or by finding spermatophore remains on filter paper). Adult females were sacrificed after their broods hatched. Adult phenotypes from treated insects were examined alongside control individuals that were injected with dsGFP. Dissections of gonads and other reproductive organs were performed in Ringer Solution. For scanning electron microscopy of tergal glands, adult insect abdomens were removed from the thorax and head, dehydrated in 100 percent ethanol, processed by critical point drying, coated with gold, and imaged on a Philips XL30 SEM microscope.

### Oligosaccharide chemistry

After dissection, tergal glands were blended with 1 ml of 80% (v/v) EtOH/H_2_0 and incubated at 4°C overnight to isolate the oligosaccharides. The supernatant was collected and dehydrated. The oligosaccharides were reconstituted in 1 M NaBH_4_.and incubated at 60°C for 2 hours. Oligosaccharides were further purified in as previously described (112). Structural elucidation employed an Agilent 6520 Q-TOF mass spectrometer paired with a 1200 series HPLC with a chip interface for structural analysis. A capillary pump was used to load and enrich the sample onto a porous graphitized carbon chip, this solvent contained 3% (v/v) ACN/H_2_O with 0.1% formic acid at a flow rate of 3 μl/min. A nano pump containing a binary solvent system was used for separation with a flow rate of 0.4 μl/min, solvent A contained 3% (v/v) ACN/H_2_O with 0.1% formic acid and solvent B contained 90% (v/v) ACN/H_2_O with 0.1% formic acid. A 45-min gradient was performed with the following conditions: 0.00-10.00 min, 2-5% B, 10.00-20.00 min, 7.5% B, 20.00-25.00 min, 10% B, 25.00-30.00 min, 99% B, 30.00-35.00 min, 99% B, 35.00-45.10, 2% B. The instrument was run in positive ion mode. Data analysis was performed with the Agilent MassHunter Qualitative Analysis (B.06) software. For quantitative determination samples were run on an Agilent 6210 TOF mass spectrometer paired with a 1200 series HPLC with a chip interface with the same chromatographic parameters.

### Behavioral assays

dsBgTra females were reared for one week on food pellets (Purina No. 5001 Rodent Diet, PMI Nutrition International) and distilled water in a temperature-regulated walk-in room at 27 ± 1°C, 40–70% relative humidity, and L:D = 12:12 photoperiod (Light, 20:00 – 8:00). A wild-type strain (wild-type, American Cyanamid strain = Orlando Normal, collected in a Florida apartment >60 yrs ago) was also kept in the same conditions. Newly emerged wild-type males and wild-type females were kept in separate cages to prevent contact. Sexually mature virgin males (20-day-old) and sexually mature virgin females (5-6 days old) were used in behavioral assays. For assays of the wing-raising display in response to isolated female and male antennae, dsBgTra females, wild-type females, and wild-type males were acclimated individually for 1 day in glass test tubes (15 cm x 2.5 cm diameter) stoppered with cotton. Observations were carried out at 17:00-19:00 h (scotophase) in the walk-in incubator room under red fluorescent lights. Each test tube was horizontally placed under an IR-sensitive camera (Everfocus EQ610 Polestar, Taiwan). The antennae of the tested insects were stimulated by contact with either an isolated wild-type female antenna or wild-type male antenna, which was attached to the tip of a glass Pasteur pipette by dental wax. Observation time was 1 min. A single antenna was used for 2-3 tested insects. The percentage of responders was compared by Chi-square test. The latency of wing-raising display was quantified in seconds, from the start of stimulation to the initiation of the display, and compared by t-test (unpaired, p < 0.05).

Observations of nuptial feeding were conducted at 12:00-19:00 hr (scotophase). As above, individual tested insects were acclimated in test tubes, but a single wild-type male or wild-type female was introduced into the test tube and wing-raising and tergal gland feeding behaviors were observed for both insects.

### Cuticular hydrocarbon analysis

To compare cuticular hydrocarbons (CHCs) between control and RNAi treatments, individual cockroaches were extracted for 5 min in 200 µL of hexane containing 10 µg of heptacosane (*n*-C26) as an internal standard. Extracts were reduced to 150 µL and 1 μl was injected in splitless mode using a 7683B Agilent autosampler into a DB-5 column (20 m × 0.18 mm internal diameter × 0.18 µm film thickness; J&W Scientific) in an Agilent 7890 series GC (Agilent Technologies) connected to a flame ionization detector with ultrahigh-purity hydrogen as carrier gas (0.75 mL/min constant flow rate). The column was held at 50 °C for 1 min, increased to 320 °C at 10 °C/min, and held at 320 °C for 10 min. For Principal Components Analysis (PCA), we used the percentage of each of 30 peaks previously identified (Jurenka et al, 1989). PCA was conducted in JMP (JMP Pro 12, SAS Institute, Inc., Cary, NC). The amount (mass) of each peak was determined relative to the *n*-C26 internal standard.

## ACKNOWLEDGEMENTS

We thank Dr. Ian Orchard (*R. prolixus*), Deanna Fox (*P. humanus*), and Dr. John Clark (*P. humanus*) for providing us with tissue samples. Megan Meany provided invaluable assistance dissecting and processing *B. germanica* tergal glands for oligosaccharide analysis. We are grateful to Dr. Carlito Lebrilla for assistance with the oligosaccharide analysis. RNA-sequencing was carried out at the DNA Technologies and Expression Analysis Cores at the UC Davis Genome Center, supported by NIH Shared Instrumentation Grant 1S10OD010786-01. J.R.W. was supported by National Science Foundation (NSF) Graduate Research Fellowship Program grant 1650042 as well as by NSF/EDEN grant number IOS # 0955517. E.K.D. was supported by NIH NRSA and UC Davis Chancellor’s fellowships. X.B. was supported by the Spanish MINECO (grants CGL2012–36251 and CGL2015–64727-P, including FEDER funds) and the Catalan Government (grant 2017 SGR 1030). C.S. and A.W-K were supported in part by NSF Division of Integrative Organismal Systems Grant Number IOS-1557864. A.K. was supported by NIH grant 5R35GM122592.

---

Supplemental Table 1: Sources of arthropod gene models used in phylogenetic analyses of the DMRT and SR gene families. Orders are color-coded as follows: holometabolous insects (orange), hemimetabolous insects (green), non-insect hexapods (red), crustaceans (blue), and chelicerates (purple).

Supplemental Table 2: Distances between *prospero* and *doublesex* in the genomes of 10 different insect species.

Supplemental Table 3: Scaffolds containing *prospero* and their sizes from nine hemipteran species.

Supplemental Table 4: Estimated percentages of cuticular hydrocarbons in wild-type and dsBgTra-treated *Blattella germanica*. Each gas chromatograph (GC) peak is represented as a percentage of the total amount of all 30 hydrocarbons. The peak numbers correspond to the CHCs identified by Jurenka et al. (1989). Peak 15 (9-, 11-, 13-, and 15-methylnonacosane) is known as a male-enriched CHC, and peak 22 (3,7-, 3,9-, and 3,11-dimethylnonacosane) is a female-enriched CHC. 3,11-Dimethylnonacosane also serves as precursor to several components of the female contact sex pheromone.

Supplemental Table 5: Primers used in RACE, RT-PCR, and qPCR in this study.

Supplemental Figure 1: Maximum likelihood gene tree of DMRT family proteins in 29 arthropod species (Supplemental Table 1) generated using RAxML (Stamatakis, 2014). Node labels indicate bootstrap support values for clades. Clades containing *Drosophila* DMRT genes are outlined in shaded boxes. *doublesex* genes that are known to regulate male and female differentiation via alternative spliceforms are indicated with asterisks (*); *doublesex* genes that promote male differentiation via transcriptional upregulation in males are indicated with a double cross (‡). Genes studied in this paper are indicated with a filled circle (●). For ease of viewing the *doublesex* clade, the tree is rooted with DMRT93B sequences clade.

Supplemental Figure 2: Transformer proteins do not form a monophyletic group in the gene tree of SR family proteins in 29 arthropod species generated with maximum likelihood (RAxML). Node labels indicate bootstrap values for clades. Previously studied *transformer* genes are indicated with asterisks; genes investigated in this paper are indicated with filled circles.

Supplemental Figure 3: MAFFT protein alignment of RpTraB and the longer female-specific isoform of RpTraA from *Rhodnius prolixus*. The CAM domain is boxed in pink, the RS domain in blue, the RRM domain in black, and the proline rich domain in green. *R. prolixus* shows the first Tra duplication reported outside of Hymenoptera (Geuverink and Beukeboom, 2014).

Supplemental Figure 4: RRM domains found in select Transformer proteins. Predicted sequences of Tra proteins from *Apis mellifera*, *Rhodnius prolixus*, *Blattella germanica*, and *Tigriopus californicus*, with the residues predicted to form an RRM domain highlighted in yellow.

Supplemental Figure 5: MAFFT protein alignment of the complete CAM domain in 9 different insect species and one crustacean. The cockroach *Blattella germanica* has three different types of transcripts: some transcripts terminate halfway through the CAM domain (*BgTra1*, *BgTra2*), others code for an intact CAM domain (*BgTra6, BgTra7, BgTra8*), and in some the inclusion of an alternatively spliced exon interrupts the CAM domain with 47 additional residues (*BgTra3, BgTra4, BgTra5*).

Supplemental Figure 6: dsBgTra reduces *BgTra* expression in males and females of *Blattella germanica*. Left panel, *BgTra* expression in the colleterial glands of females injected with dsTra or dsGFP (control) throughout development (n=2 dsTra and n=2 dsGFP). Middle panel, *BgTra* expression in the accessory glands of males injected with dsTra and dsGFP (control) throughout development (n=3 dsTra and n=3 dsGFP). Right panel, *BgTra* in the fat body of females injected with dsTra after terminal molt (n=3 dsTra and n=3 dsGFP). This last group of females were used to study the maternal effects of *BgTra* on embryonic development. Primers used for qPCR span junction between exons 5 and 6, which is shared by all *BgTra* isoforms. All graphs show gene expression measured by qPCR, relative to β-*actin*.

Supplemental Figure 7: Abnormal morphology in 6^th^ instar dsBgTra female nymphs of *Blattella germanica*. Tissue extrusion in dsBgTra females (left) compared with the normal abdomen in wild-type 6^th^ instar females (right).

Supplemental Figure 8: *BgTra* controls sex-specific oligosaccharide synthesis in tergal glands (A) Relative abundance of oligosaccharides in tergal glands derived from dsBgTra females (dsTra1-dsTra10) and wild-type males (wt1-wt10). Each bar represents one individual, with different compounds shown in different colors. (B) Principal components analysis on the relative proportions of different oligosaccharides in dsBgTra female and wild-type male tergal glands. These two groups sort into different parts of PCA space, indicating that although all sugars produced by wild type male tergal glands are present in tergal glands from dsBgTra females, they produce these sugars in different relative amounts. Dimension 1 is primarily driven by the relative abundance of a non-reducing hexose with mass 990.328.

Supplemental Figure 9: *BgTra* controls production of sex-specific cuticular hydrocarbons. Gas chromatograms (A) and PCA (B, C) of the cuticular hydrocarbons of *Blattella germanica*. The peak numbers correspond to the CHCs in Supplemental Table 4 and to Jurenka et al. (1989). In the score plot (B), the solid black squares represent the centroids of their respective groups. Wild-type males in solid blue circles (n= 10), wild-type females in open red circles (n=9), dsBgTra females in green diamonds (n=9). In the loadings plot (C), note that peak 15, which is abundant in wild-type males, pulls towards wild-type males and dsBgTra females (to the left), whereas peak 22, which is more abundant in females, pulls towards wild-type females (to the right). (D, E) Statistical analysis of the percentage representation of peak 15 (D) and peak 22 (E) in the total cuticular hydrocarbons of wild-type males, wild-type females and dsBgTra females. Peak 15 (9-, 11-, 13-, and 15-methylnonacosane), which is more common in wild-type males, was more represented in dsBgTra females than in wild-type females. Peak 22 (3,7-, 3,9-, and 3,11-dimethylnonacosane), which serves as precursor to several components of the female contact sex pheromone, was significantly under-represented in dsBgTra females compared to wild-type females. (F) Statistical analysis of the total cuticular hydrocarbons of wild-type males, wild-type females and dsBgTra females. In the box plots, the horizontal line within the box represents the median value and the box represents the 25^th^ to 75^th^ quantiles. Letters within each graph represent the results of Tukey’s HSD, and different letters represent statistically significant differences (*P* < 0.05).

Supplemental Figure 10: *BgDsx* RNAi causes an increase in *BgDsx* transcript abundance in both males and females of *Blattella germanica*. *BgDsx* expression was measured (A) 6 h and (B) 3 days after injection with dsRNA targeting the *BgDsx* common region. (C) qPCR primers (red arrows) were located outside the region targeted by dsBgDsx (purple bar). n= 3 females and 3 males per treatment. Expression levels of *BgDsx* are normalized to β-*actin*.

Supplemental Figure 11: Representative sampling of tissue extrusions found in dsBgDsx males of *Blattella germanica* (light blue arrowheads).

Supplemental Figure 12: The conglobate glands and utricles of dsBgDsx males of *Blattella germanica* fail to properly mature. (A) Utricles and seminal vesicles from five day old dsBgDsx male. Note tissue discoloration of utricles (light blue arrows). (B) Utricles and seminal vesicles of untreated control five day old male. (C) Conglobate gland from five day old dsBgDsx male is malformed compared to control conglobate gland from untreated five day old male (D).

Supplemental Figure 13: BgDsx has no effect on brood size in females of *Blattella germanica* that were injected with dsBgDsx either at multiple stages throughout nymphal development (A) or as virgin adult females (B).

Supplemental File 1: Queries used in BLASTP searches of arthropod gene models for DMRT homologs.

Supplemental File 2: Queries used in BLASTP searches of arthropod gene models for Transformer and SR family proteins.

## REFERENCES

1. Charlesworth B. The evolution of chromosomal sex determination and dosage compensation. Curr Biol. 1996 Feb 1;6(2):149–62.

2. Graves JAM. Sex chromosome specialization and degeneration in mammals. Cell. 2006 Mar 10;124(5):901–14.

3. Ellegren H. Evolution of the avian sex chromosomes and their role in sex determination. Trends Ecol Evol. 2000 May 1;15(5):188–92.

4. Jablonka E, Lamb MJ. The evolution of heteromorphic sex chromosomes. Biol Rev. 1990 Aug 1;65(3):249–76.

5. Ser JR, Roberts RB, Kocher TD. Multiple interacting loci control sex determination in Lake Malawi cichlid fish. Evolution. 2010 Feb 1;64(2):486–501.

6. Beye M, Hasselmann M, Fondrk MK, Page RE, Omholt SW. The gene csd is the primary signal for sexual development in the honeybee and encodes an SR-type protein. Cell. 2003 Aug 22;114(4):419–29.

7. Merchant-Larios H, Díaz-Hernández V. Environmental sex determination mechanisms in reptiles. Sex Dev. 2013;7(1–3):95–103.

8. Bewick AJ, Anderson DW, Evans BJ. Evolution of the closely related, sex-related genes *Dm-W* and *Dmrt1* in African clawed frogs (*Xenopus*). Evolution. 2011;65(3):698–712.

9. Roco ÁS, Olmstead AW, Degitz SJ, Amano T, Zimmerman LB, Bullejos M. Coexistence of Y, W, and Z sex chromosomes in *Xenopus tropicalis*. Proc Natl Acad Sci. 2015 Aug 25;112(34):E4752–61.

10. Myosho T, Otake H, Masuyama H, Matsuda M, Kuroki Y, Fujiyama A, et al. Tracing the emergence of a novel sex-determining gene in Medaka, *Oryzias luzonensis*. Genetics. 2012 May;191(1):163–70.

11. Takehana Y, Hamaguchi S, Sakaizumi M. Different origins of ZZ/ZW sex chromosomes in closely related medaka fishes, Oryzias javanicus and O. hubbsi. Chromosome Res. 2008 Aug 1;16(5):801–11.

12. Li J, Phillips RB, Harwood AS, Koop BF, Davidson WS. Identification of the sex chromosomes of brown trout (*Salmo trutta*) and their comparison with the corresponding chromosomes in Atlantic salmon (*Salmo salar*) and rainbow trout (*Oncorhynchus mykiss*). Cytogenet Genome Res. 2011;133(1):25–33.

13. Bachtrog D, Mank JE, Peichel CL, Kirkpatrick M, Otto SP, Ashman T-L, et al. Sex determination: why so many ways of doing it? PLOS Biol. 2014 Jul 1;12(7):e1001899.

14. Anderson JL, Mari AR, Braasch I, Amores A, Hohenlohe P, Batzel P, et al. Multiple sex-associated regions and a putative sex chromosome in zebrafish revealed by RAD mapping and population genomics. Plos One. 2012 Jul 9;7(7):e40701.

15. Liew WC, Bartfai R, Lim Z, Sreenivasan R, Siegfried KR, Orban L. Polygenic sex determination system in zebrafish. Plos One. 2012 Apr 10;7(4):e34397.

16. Tomita T, Wada Y. Multifactorial sex determination in natural-populations of the housefly (*Musca domestica*) in Japan. Jpn J Genet. 1989 Oct;64(5):373–82.

17. Meisel RP, Davey T, Son JH, Gerry AC, Shono T, Scott JG. Is multifactorial sex determination in the house fly, Musca domestica (L.), stable over time? J Hered. 2016 Jan 1;107(7):615–25.

18. Nakamura M. Sex determination in amphibians. Semin Cell Dev Biol. 2009 May 1;20(3):271–82.

19. Duboule D. Temporal colinearity and the phylotypic progression: a basis for the stability of a vertebrate Bauplan and the evolution of morphologies through heterochrony. Dev Camb Engl Suppl. 1994;135–42.

20. Raff RA. The shape of life: genes, development, and the evolution of animal form. University of Chicago Press; 2012. 545 p.

21. Hazkani-Covo E, Wool D, Graur D. In search of the vertebrate phylotypic stage: A molecular examination of the developmental hourglass model and von Baer’s third law. J Exp Zoolog B Mol Dev Evol. 2005;304B(2):150–8.

22. Raymond CS, Kettlewell JR, Hirsch B, Bardwell VJ, Zarkower D. Expression of *Dmrt1* in the genital ridge of mouse and chicken embryos suggests a role in vertebrate sexual development. Dev Biol. 1999 Nov 15;215(2):208–20.

23. Brennan J, Capel B. One tissue, two fates: molecular genetic events that underlie testis versus ovary development. Nat Rev Genet. 2004 Jul;5(7):509–21.

24. Hau M. Regulation of male traits by testosterone: implications for the evolution of vertebrate life histories. BioEssays. 2007 Feb 1;29(2):133–44.

25. Maatouk DM, DiNapoli L, Alvers A, Parker KL, Taketo MM, Capel B. Stabilization of *beta-catenin* in XY gonads causes male-to-female sex-reversal. Hum Mol Genet. 2008 Oct 1;17(19):2949–55.

26. Matson CK, Murphy MW, Sarver AL, Griswold MD, Bardwell VJ, Zarkower D. *DMRT1* prevents female reprogramming in the postnatal mammalian testis. Nature. 2011 Aug;476(7358):101.

27. Lindeman RE, Gearhart MD, Minkina A, Krentz AD, Bardwell VJ, Zarkower D. Sexual cell-fate reprogramming in the ovary by *DMRT1*. Curr Biol. 2015 Mar 16;25(6):764–71.

28. Herpin A, Schartl M. Plasticity of gene-regulatory networks controlling sex determination: of masters, slaves, usual suspects, newcomers, and usurpators. EMBO Rep. 2015 Oct 1;16(10):1260–74.

29. Pires-daSilva A, Sommer RJ. Conservation of the global sex determination gene tra-1 in distantly related nematodes. Genes Dev. 2004 May 15;18(10):1198–208.

30. Ohbayashi F, Suzuki MG, Mita K, Okano K, Shimada T. A homologue of the *Drosophila doublesex* gene is transcribed into sex-specific mRNA isoforms in the silkworm, *Bombyx mori*. Comp Biochem Physiol B Biochem Mol Biol. 2001 Jan 1;128(1):145–58.

31. Suzuki MG, Funaguma S, Kanda T, Tamura T, Shimada T. Analysis of the biological functions of a doublesex homologue in *Bombyx mori*. Dev Genes Evol. 2003 Jul 1;213(7):345–54.

32. Cho S, Huang ZY, Zhang J. Sex-specific splicing of the honeybee *doublesex* gene reveals 300 million years of evolution at the bottom of the insect sex-determination pathway. Genetics. 2007 Nov 1;177(3):1733–41.

33. Hasselmann M, Gempe T, Schiøtt M, Nunes-Silva CG, Otte M, Beye M. Evidence for the evolutionary nascence of a novel sex determination pathway in honeybees. Nature. 2008 Jul;454(7203):519–22.

34. Oliveira DCSG, Werren JH, Verhulst EC, Giebel JD, Kamping A, Beukeboom LW, et al. Identification and characterization of the *doublesex* gene of *Nasonia*. Insect Mol Biol. 2009;18(3):315–24.

35. Saccone G, Pane A, Polito LC. Sex determination in flies, fruitflies and butterflies. Genetica. 2002 Sep 1;116(1):15–23.

36. Shukla JN, Palli SR. Sex determination in beetles: production of all male progeny by parental RNAi knockdown of *transformer*. Sci Rep. 2012 Aug 24;2:602.

37. Shukla JN, Palli SR. *Doublesex* target genes in the red flour beetle, *Tribolium castaneum*. Sci Rep. 2012 Dec 10;2:948.

38. Barnes TM, Hodgkin J. The *tra-3* sex determination gene of *Caenorhabditis elegans* encodes a member of the calpain regulatory protease family. Embo J. 1996 Sep 2;15(17):4477–84.

39. Pilgrim D, Mcgregor A, Jackle P, Johnson T, Hansen D. The *C. elegans* sex-determining gene *fem-2* encodes a putative protein phosphatase. Mol Biol Cell. 1995 Sep;6(9):1159– 71.

40. Pires-daSilva A. Evolution of the control of sexual identity in nematodes. Semin Cell Dev Biol. 2007 Jun 1;18(3):362–70.

41. Berkseth M, Ikegami K, Arur S, Lieb JD, Zarkower D. TRA-1 ChIP-seq reveals regulators of sexual differentiation and multilevel feedback in nematode sex determination. Proc Natl Acad Sci U S A. 2013 Oct 1;110(40):16033–8.

42. Kopp A. *Dmrt* genes in the development and evolution of sexual dimorphism. Trends Genet. 2012 Apr 1;28(4):175–84.

43. Matson CK, Zarkower D. Sex and the singular DM domain: insights into sexual regulation, evolution and plasticity. Nat Rev Genet. 2012 Mar;13(3):163–74.

44. Smith CA, Katz M, Sinclair AH. *DMRT1* is upregulated in the gonads during female-to-male sex reversal in ZW chicken embryos. Biol Reprod. 2003 Feb;68(2):560–70.

45. Webster KA, Schach U, Ordaz A, Steinfeld JS, Draper BW, Siegfried KR. *Dmrt1* is necessary for male sexual development in zebrafish. Dev Biol. 2017 Feb 1;422(1):33–46.

46. Shen MM, Hodgkin J. *mab-3*, a gene required for sex-specific yolk protein expression and a male-specific lineage in *C. elegans*. Cell. 1988 Sep 23;54(7):1019–31.

47. Portman DS. Genetic control of sex differences in C. elegans neurobiology and behavior. In: Yamamoto D, editor. Genetics of sexual differentiation and sexually dimorphic behaviors. San Diego: Elsevier Academic Press Inc; 2007. p. 1–37.

48. Mason DA, Rabinowitz JS, Portman DS. *dmd-3*, a *doublesex*-related gene regulated by *tra-1*, governs sex-specific morphogenesis in *C. elegans*. Development. 2008 Jul 15;135(14):2373–82.

49. Siehr MS, Koo PK, Sherlekar AL, Bian X, Bunkers MR, Miller RM, et al. Multiple *doublesex*-related genes specify critical cell fates in a *C. elegans* male neural circuit. Plos One. 2011 Nov 1;6(11):e26811.

50. Serrano-Saiz E, Oren-Suissa M, Bayer EA, Hobert O. Sexually dimorphic differentiation of a *C. elegans* hub neuron is cell autonomously controlled by a conserved transcription factor. Curr Biol. 2017 Jan 23;27(2):199–209.

51. Baker BS, Ridge KA. Sex and the single cell. I. On the action of major loci affecting sex determination in *Drosophila melanogaster*. Genetics. 1980 Feb 1;94(2):383–423.

52. Rideout EJ, Dornan AJ, Neville MC, Eadie S, Goodwin SF. Control of sexual differentiation and behavior by the *doublesex* gene in *Drosophila melanogaster*. Nat Neurosci. 2010 Apr;13(4):458–66.

53. Kato Y, Kobayashi K, Watanabe H, Iguchi T. Environmental sex determination in the Branchiopod crustacean *Daphnia magna*: deep conservation of a *doublesex* gene in the sex-determining pathway. PLOS Genet. 2011 Mar 24;7(3):e1001345.

54. Pomerantz AF, Hoy MA. Expression analysis of *Drosophila doublesex*, *transformer-2*, *intersex*, *fruitless-like*, and *vitellogenin* homologs in the parahaploid predator *Metaseiulus occidentalis* (Chelicerata: Acari: Phytoseiidae). Exp Appl Acarol. 2015 Jan; 65(1):1–16.

55. Burtis KC, Baker BS. *Drosophila doublesex* gene controls somatic sexual differentiation by producing alternatively spliced mRNAs encoding related sex-specific polypeptides. Cell. 1989 Mar 24;56(6):997–1010.

56. Wexler JR, Plachetzki DC, Kopp A. Pan-metazoan phylogeny of the *DMRT* gene family: a framework for functional studies. Dev Genes Evol. 2014 Jun;224(3):175–81.

57. Li S, Li F, Yu K, Xiang J. Identification and characterization of a *doublesex* gene which regulates the expression of insulin-like androgenic gland hormone in *Fenneropenaeus chinensis*. Gene. 2018 Apr 5;649:1–7.

58. Zhuo J-C, Hu Q-L, Zhang H-H, Zhang M-Q, Jo SB, Zhang C-X. Identification and functional analysis of the *doublesex* gene in the sexual development of a hemimetabolous insect, the brown planthopper. Insect Biochem Mol Biol. 2018 Nov 1;102:31–42.

59. Long JC, Caceres JF. The SR protein family of splicing factors: master regulators of gene expression. Biochem J. 2009 Jan 1;417(1):15–27.

60. Verhulst EC, van de Zande L, Beukeboom LW. Insect sex determination: it all evolves around *transformer*. Curr Opin Genet Dev. 2010 Aug;20(4):376–83.

61. Hediger M, Henggeler C, Meier N, Perez R, Saccone G, Bopp D. Molecular characterization of the key switch F provides a basis for understanding the rapid divergence of the sex-determining pathway in the housefly. Genetics. 2010 Jan;184(1):155–U280.

62. Tanaka A, Aoki F, Suzuki MG. Conserved domains in the *transformer* protein act complementary to regulate sex-specific splicing of its own pre-mRNA. Sex Dev. 2018;12(4):180–90.

63. Boggs RT, Gregor P, Idriss S, Belote JM, McKeown M. Regulation of sexual differentiation in *D. melanogaster* via alternative splicing of RNA from the *transformer* gene. Cell. 1987 Aug 28;50(5):739–47.

64. Sosnowski BA, Belote JM, McKeown M. Sex-specific alternative splicing of RNA from the *transformer* gene results from sequence-dependent splice site blockage. Cell. 1989 Aug 11;58(3):449–59.

65. Inoue K, Hoshijima K, Higuchi I, Sakamoto H, Shimura Y. Binding of the *Drosophila transformer* and *transformer-2* proteins to the regulatory elements of *doublesex* primary transcript for sex-specific RNA processing. Proc Natl Acad Sci U S A. 1992 Sep 1;89(17):8092–6.

66. Mesquita RD, Vionette-Amaral RJ, Lowenberger C, Rivera-Pomar R, Monteiro FA, Minx P, et al. Genome of *Rhodnius prolixus*, an insect vector of Chagas disease, reveals unique adaptations to hematophagy and parasite infection. Proc Natl Acad Sci. 2015 Dec 1;112(48):14936–41.

67. Marchler-Bauer A, Bo Y, Han L, He J, Lanczycki CJ, Lu S, et al. CDD/SPARCLE: functional classification of proteins via subfamily domain architectures. Nucleic Acids Res. 2017 Jan 4;45(D1):D200–3.

68. Hoshijima K, Inoue K, Higuchi I, Sakamoto H, Shimura Y. Control of *doublesex* alternative splicing by *transformer* and *transformer-2* in *Drosophila*. Science. 1991 May 10;252(5007):833–6.

69. Geuverink E, Beukeboom LW. Phylogenetic distribution and evolutionary dynamics of the sex determination genes *doublesex* and *transformer* in insects. Sex Dev. 2014;8(1– 3):38–49.

70. Grabherr MG, Haas BJ, Yassour M, Levin JZ, Thompson DA, Amit I, et al. Full-length transcriptome assembly from RNA-seq data without a reference genome. Nat Biotechnol. 2011 Jul;29(7):644–52.

71. Mitchell AL, Attwood TK, Babbitt PC, Blum M, Bork P, Bridge A, et al. InterPro in 2019: improving coverage, classification and access to protein sequence annotations. Nucleic Acids Res. 2019 Jan 8;47(D1):D351–60.

72. Kijimoto T, Moczek AP, Andrews J. Diversification of *doublesex* function underlies morph-, sex-, and species-specific development of beetle horns. Proc Natl Acad Sci. 2012 Dec 11;109(50):20526–31.

73. Ruiz MF, Stefani RN, Mascarenhas RO, Perondini ALP, Selivon D, Sánchez L. The gene *doublesex* of the fruit fly *Anastrepha obliqua* (Diptera, Tephritidae). Genetics. 2005 Oct 1;171(2):849–54.

74. Nojima S, Sakuma M, Nishida R, Kuwahara Y. A glandular gift in the German cockroach, *Blattella germanica* (L.) (Dictyoptera : Blattellidae): The courtship feeding of a female on secretions from male tergal glands. J Insect Behav. 1999 Sep;12(5):627–40.

75. Belles X, Ylla G. Towards understanding the molecular basis of cockroach tergal gland morphogenesis. A transcriptomic approach. Insect Biochem Mol Biol. 2015 Aug 1;63:104–12.

76. Ferveur JF, Savarit F, OKane CJ, Sureau G, Greenspan RJ, Jallon JM. Genetic feminization of pheromones and its behavioral consequences in *Drosophila* males. Science. 1997 Jun 6;276(5318):1555–8.

77. Shirangi TR, Dufour HD, Williams TM, Carroll SB. Rapid evolution of sex pheromone-producing enzyme expression in *Drosophila*. PLOS Biol. 2009 Aug 4;7(8):e1000168.

78. Fan Y, Eliyahu D, Schal C. Cuticular hydrocarbons as maternal provisions in embryos and nymphs of the cockroach *Blattella germanica*. J Exp Biol. 2008 Feb 15;211(4):548–54.

79. Schal C, Gu X, Burns EL, Blomquist GJ. Patterns of biosynthesis and accumulation of hydrocarbons and contact sex pheromone in the female german cockroach, *Blattella germanica*. Arch Insect Biochem Physiol. 1994 Jan 1;25(4):375–91.

80. Ryner LC, Goodwin SF, Castrillon DH, Anand A, Villella A, Baker BS, et al. Control of male sexual behavior and sexual orientation in *Drosophila* by the *fruitless* gene. Cell. 1996 Dec 13;87(6):1079–89.

81. Heinrichs V, Ryner LC, Baker BS. Regulation of sex-specific selection of *fruitless* 5′ splice sites by *transformer* and *transformer-2*. Mol Cell Biol. 1998 Jan;18(1):450–8.

82. Kimura K-I, Ote M, Tazawa T, Yamamoto D. *Fruitless* specifies sexually dimorphic neural circuitry in the *Drosophila* brain. Nature. 2005 Nov 10;438(7065):229–33.

83. Meier N, Käppeli SC, Niessen MH, Billeter J-C, Goodwin SF, Bopp D. Genetic control of courtship behavior in the housefly: evidence for a conserved bifurcation of the sex-determining pathway. PLOS ONE. 2013 Apr 22;8(4):e62476.

84. Clynen E, Ciudad L, Bellés X, Piulachs M-D. Conservation of *fruitless* role as master regulator of male courtship behaviour from cockroaches to flies. Dev Genes Evol. 2011 May 1;221(1):43–8.

85. Salvemini M, Polito C, Saccone G. Fruitless alternative splicing and sex behaviour in insects: an ancient and unforgettable love story? J Genet. 2010;89(3):287–299.

86. Roth LM, Willis ER. A study of cockroach behavior. Am Midl Nat. 1952;47(1):66–129.

87. Stevens NM. Studies in spermatogenesis: with especial reference to the “accessory chromosome.” Carnegie Institution of Washington; 1905. 114 p.

88. Nakamura T, Extavour CG. The transcriptional repressor *Blimp-1* acts downstream of BMP signaling to generate primordial germ cells in the cricket *Gryllus bimaculatus*. Development. 2016 Jan 15;143(2):255–63.

89. Xiang J, Reding K, Heffer A, Pick L. Conservation and variation in pair-rule gene expression and function in the intermediate-germ beetle *Dermestes maculatus*. Development. 2017 Dec 15;144(24):4625–36.

90. Hales KG, Fuller MT. Developmentally regulated mitochondrial fusion mediated by a conserved, novel, predicted GTPase. Cell. 1997 Jul 11;90(1):121–9.

91. Mozdy AD, Shaw JM. A fuzzy mitochondrial fusion apparatus comes into focus. Nat Rev Mol Cell Biol. 2003 Jun;4(6):468–78.

92. Endo K, Akiyama T, Kobayashi S, Okada M. *degenerative spermatocyte*, a novel gene encoding a transmembrane protein required for the initiation of meiosis in *Drosophila spermatogenesis*. Mol Gen Genet MGG. 1996 Nov 1;253(1–2):157–65.

93. Endo, K, Matsuda, Y, and Kobayashi, S. *Mdes*, a homolog of the *Drosophila degenerative spermatocyte* gene, is expressed during mouse spermatogenesis. Dev Growth Differ. 1997 Jun;(3):399–403.

94. Coschigano KT, Wensink PC. Sex-specific transcriptional regulation by the male and female *doublesex* proteins of *Drosophila*. Genes Dev. 1993 Jan 1;7(1):42–54.

95. Guo L, Xie W, Liu Y, Yang Z, Yang X, Xia J, et al. Identification and characterization of *doublesex* in *Bemisia tabaci*. Insect Mol Biol. 2018;27(5):620–32.

96. Kiuchi T, Koga H, Kawamoto M, Shoji K, Sakai H, Arai Y, et al. A single female-specific piRNA is the primary determiner of sex in the silkworm. Nature. 2014 May 29;509(7502):633–6.

97. Nagoshi RN, McKeown M, Burtis KC, Belote JM, Baker BS. The control of alternative splicing at genes regulating sexual differentiation in *D. melanogaster*. Cell. 1988 Apr 22;53(2):229–36.

98. Sasaki G, Ishiwata K, Machida R, Miyata T, Su Z-H. Molecular phylogenetic analyses support the monophyly of Hexapoda and suggest the paraphyly of Entognatha. BMC Evol Biol. 2013 Oct 31;13(1):236.

99. Ishiwata K, Sasaki G, Ogawa J, Miyata T, Su Z-H. Phylogenetic relationships among insect orders based on three nuclear protein-coding gene sequences. Mol Phylogenet Evol. 2011 Feb;58(2):169–80.

100. Misof B, Liu S, Meusemann K, Peters RS, Donath A, Mayer C, et al. Phylogenomics resolves the timing and pattern of insect evolution. Science. 2014 Nov 7;346(6210):763–7.

101. Johnson KP, Dietrich CH, Friedrich F, Beutel RG, Wipfler B, Peters RS, et al. Phylogenomics and the evolution of hemipteroid insects. Proc Natl Acad Sci U S A. 2018 Dec 11;115(50):12775–80.

102. Freitas L, Mello B, Schrago CG. Multispecies coalescent analysis confirms standing phylogenetic instability in Hexapoda. J Evol Biol. 2018 Nov;31(11):1623–31.

103. Li H, Shao R, Song N, Song F, Jiang P, Li Z, et al. Higher-level phylogeny of paraneopteran insects inferred from mitochondrial genome sequences. Sci Rep. 2015 Feb 23;5:8527.

104. Katoh K, Rozewicki J, Yamada KD. MAFFT online service: multiple sequence alignment, interactive sequence choice and visualization. Brief Bioinform [Internet]. [cited 2018 May 6]; Available from: https://academic.oup.com/bib/advance-article/doi/10.1093/bib/bbx108/4106928

105. Capella-Gutiérrez S, Silla-Martínez JM, Gabaldón T. trimAl: a tool for automated alignment trimming in large-scale phylogenetic analyses. Bioinformatics. 2009 Aug 1;25(15):1972–3.

106. Darriba D, Taboada GL, Doallo R, Posada D. ProtTest 3: fast selection of best-fit models of protein evolution. Bioinforma Oxf Engl. 2011 Apr 15;27(8):1164–5.

107. Stamatakis A. RAxML version 8: a tool for phylogenetic analysis and post-analysis of large phylogenies. Bioinformatics. 2014 May 1;30(9):1312–3.

108. Bolger AM, Lohse M, Usadel B. Trimmomatic: a flexible trimmer for Illumina sequence data. Bioinforma Oxf Engl. 2014 Aug 1;30(15):2114–20.

109. Haas BJ, Papanicolaou A, Yassour M, Grabherr M, Blood PD, Bowden J, et al. *De novo* transcript sequence reconstruction from RNA-seq using the Trinity platform for reference generation and analysis. Nat Protoc. 2013 Aug;8(8):1494–512.

110. Schmittgen TD, Livak KJ. Analyzing real-time PCR data by the comparative *C_T_* method. Nat Protoc. 2008 Jun;3(6):1101–8.

111. Katoh K, Misawa K, Kuma K, Miyata T. MAFFT: a novel method for rapid multiple sequence alignment based on fast Fourier transform. Nucleic Acids Res. 2002 Jul 15;30(14):3059–66.

112. Ninonuevo MR, Lebrilla CB. Mass spectrometric methods for analysis of oligosaccharides in human milk. Nutr Rev. 2009 Nov;67(11):S216–26.

