## Supplementary figures and images for "Hemimetabolous insects elucidate the origin of sexual development via alternative splicing"

### Supplemental Data 1

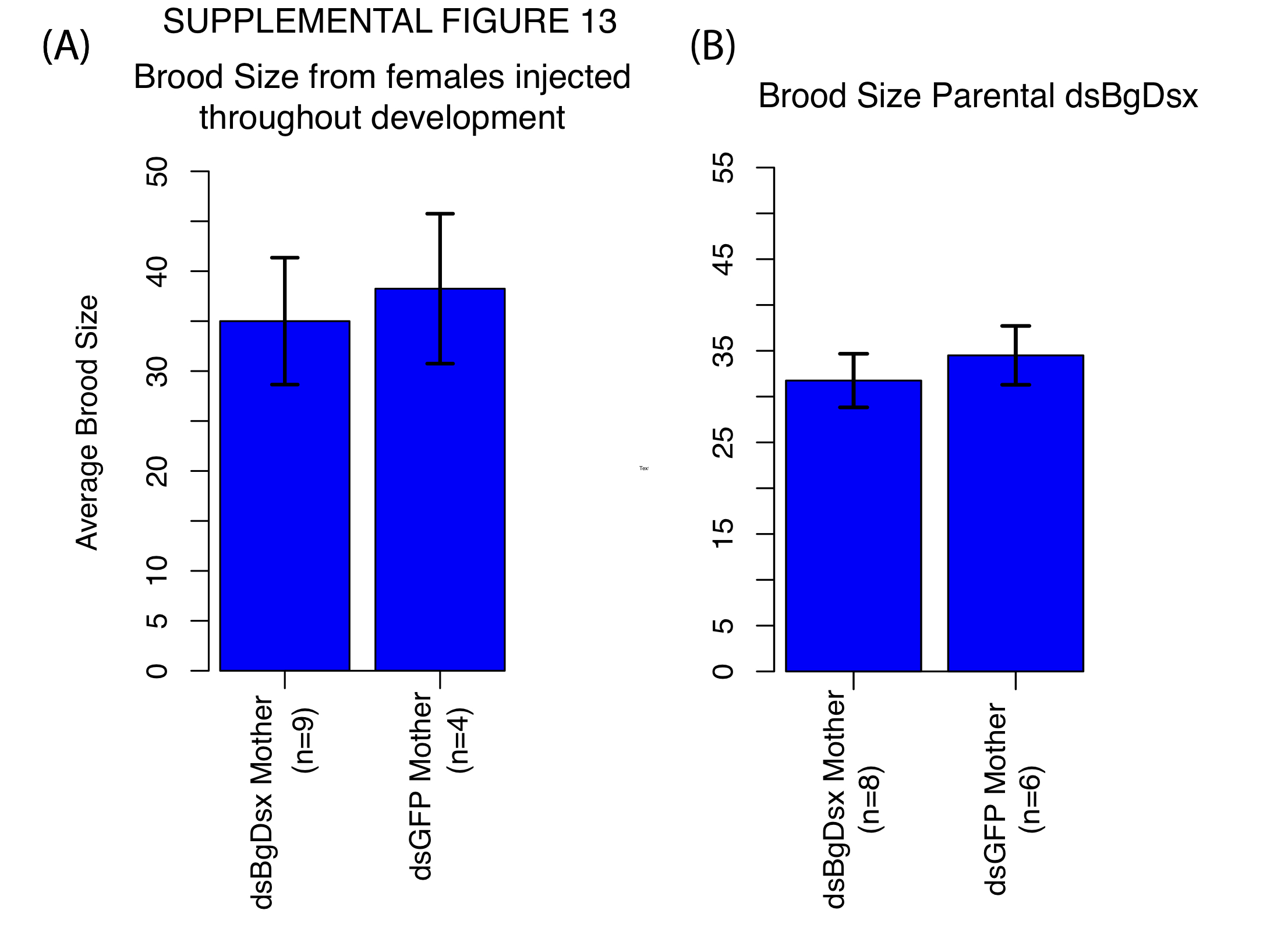

### Supplemental Figure 1

#### SUPPLEMENTAL FIGURE 1

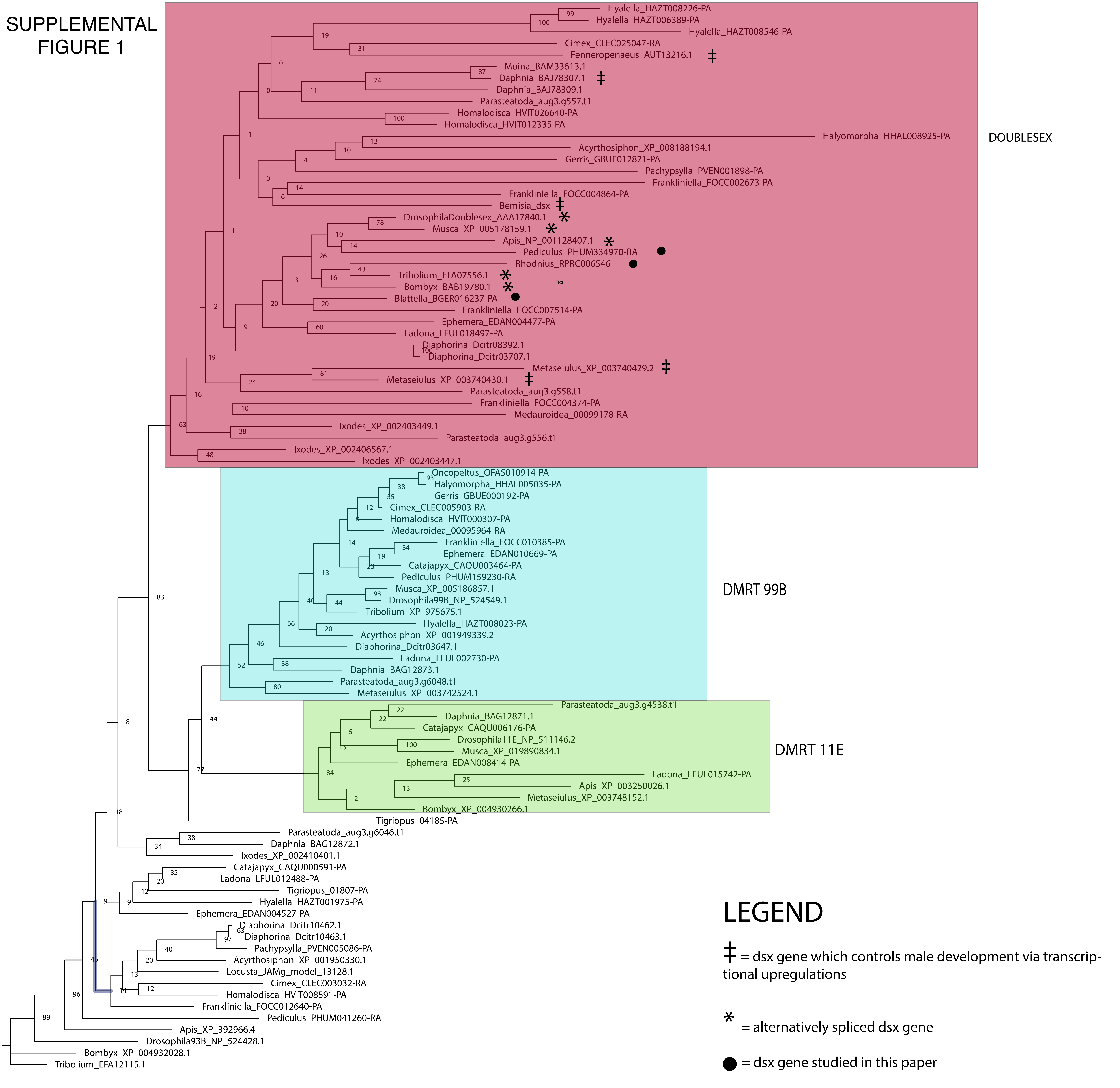

### Supplemental Figure 2

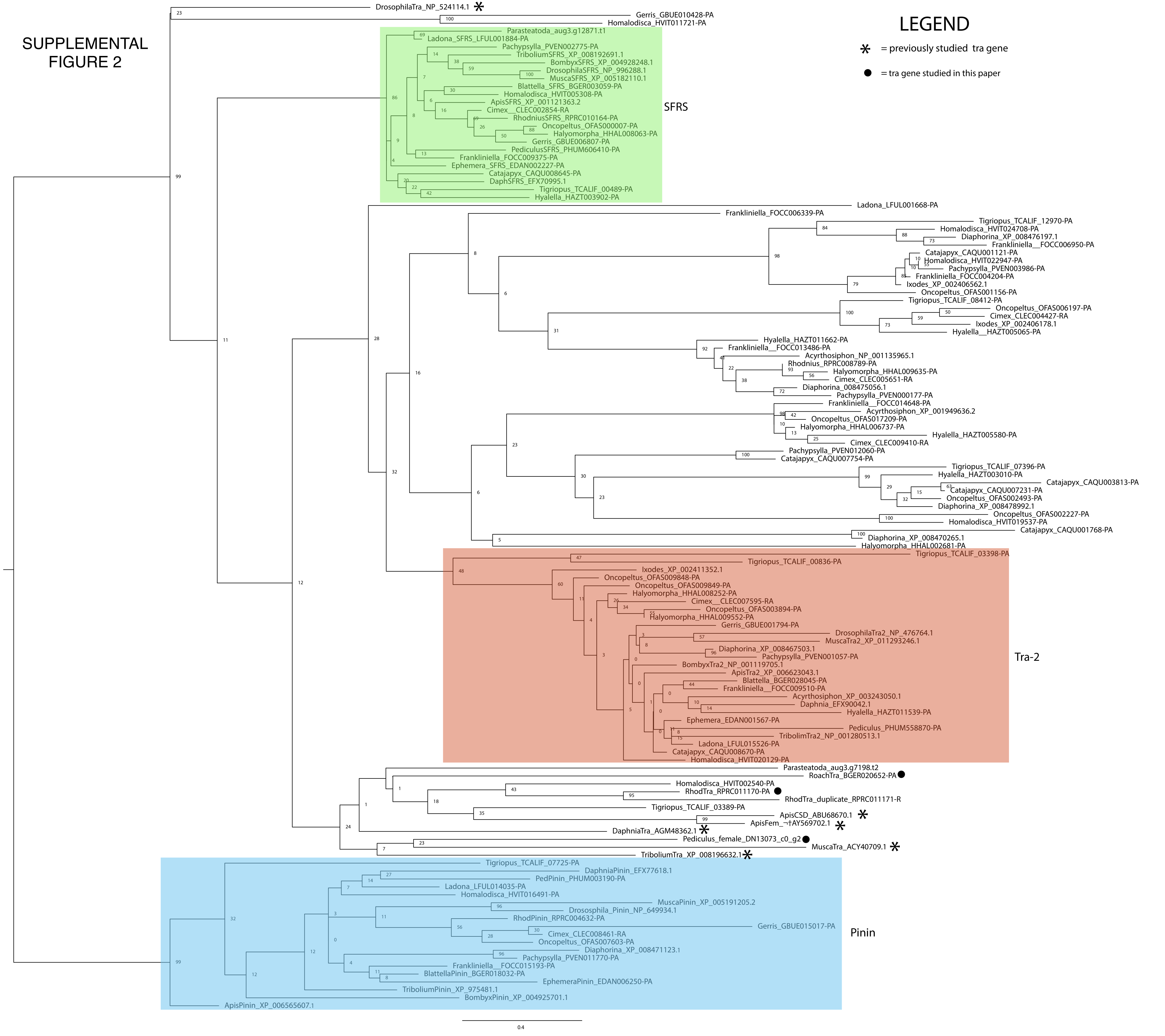

### Supplemental Figure 3

#### SUPPLEMENTAL FIGURE 3

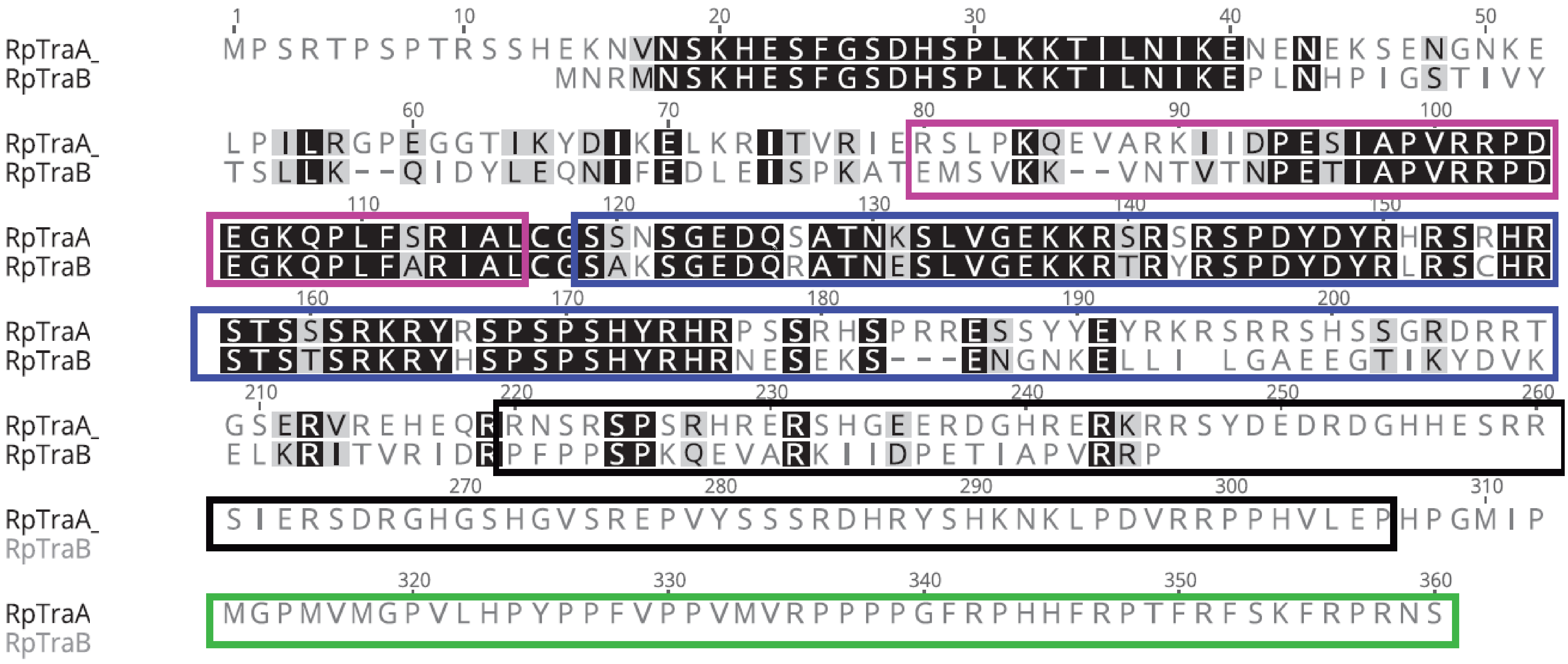

### Supplemental Figure 4

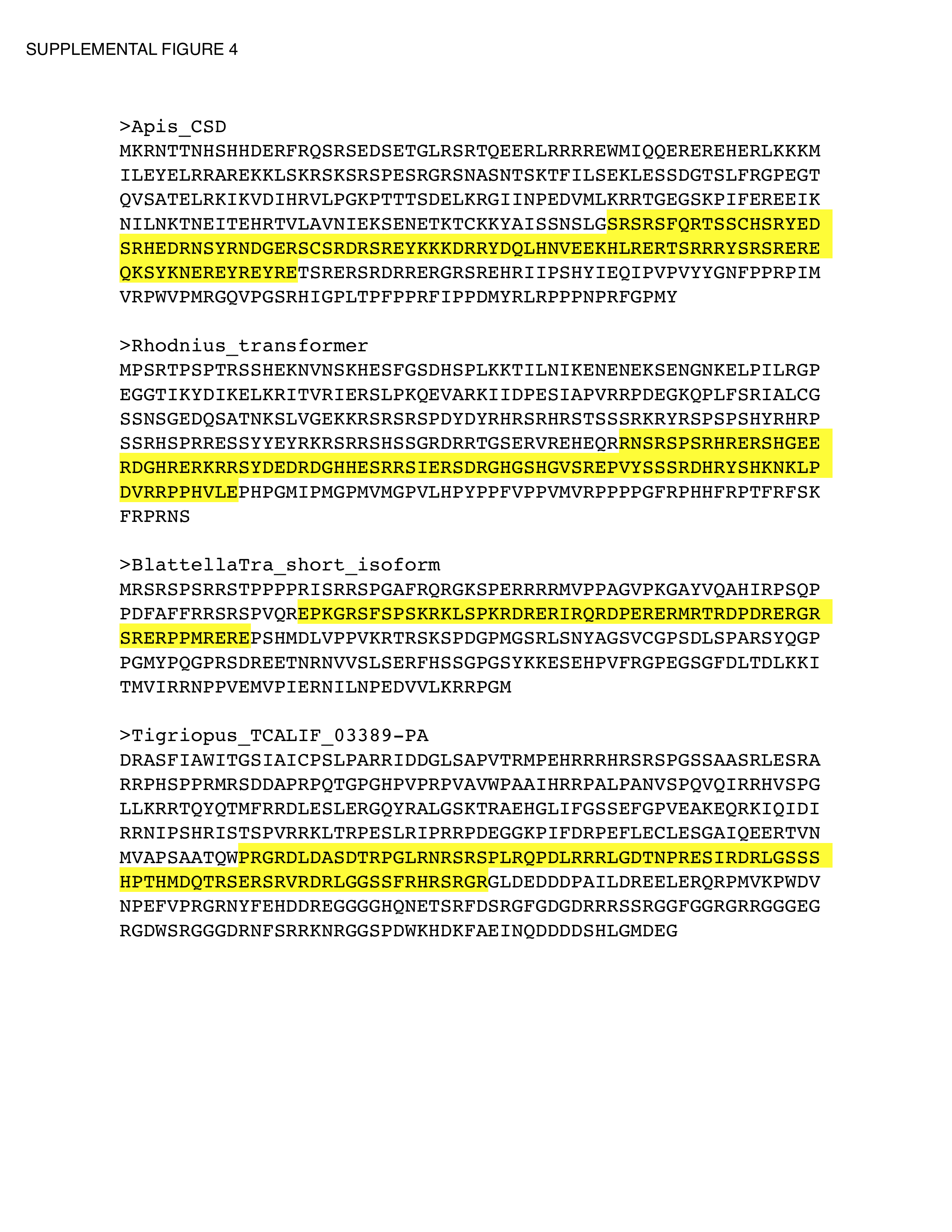

### Supplemental Figure 5

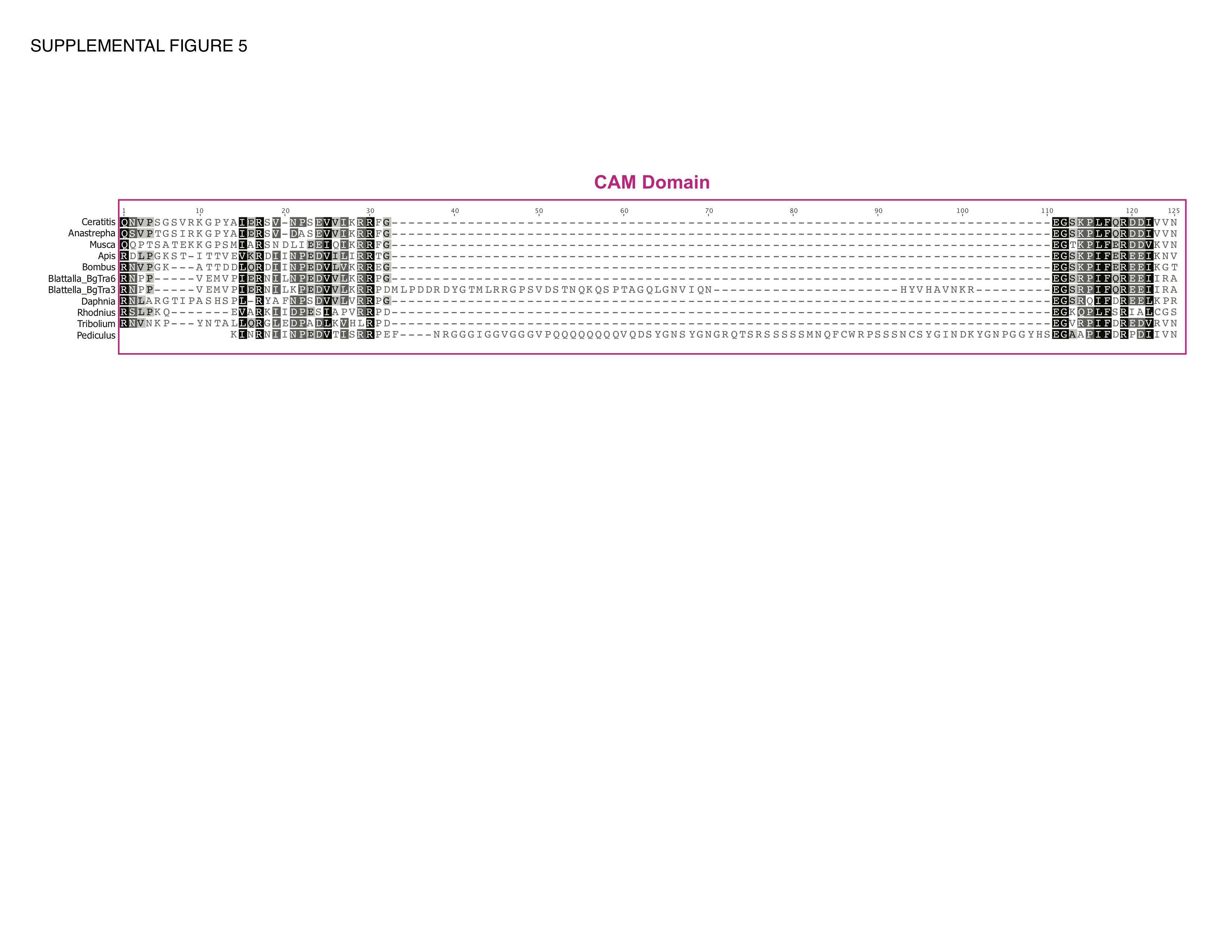

### Supplemental Figure 6

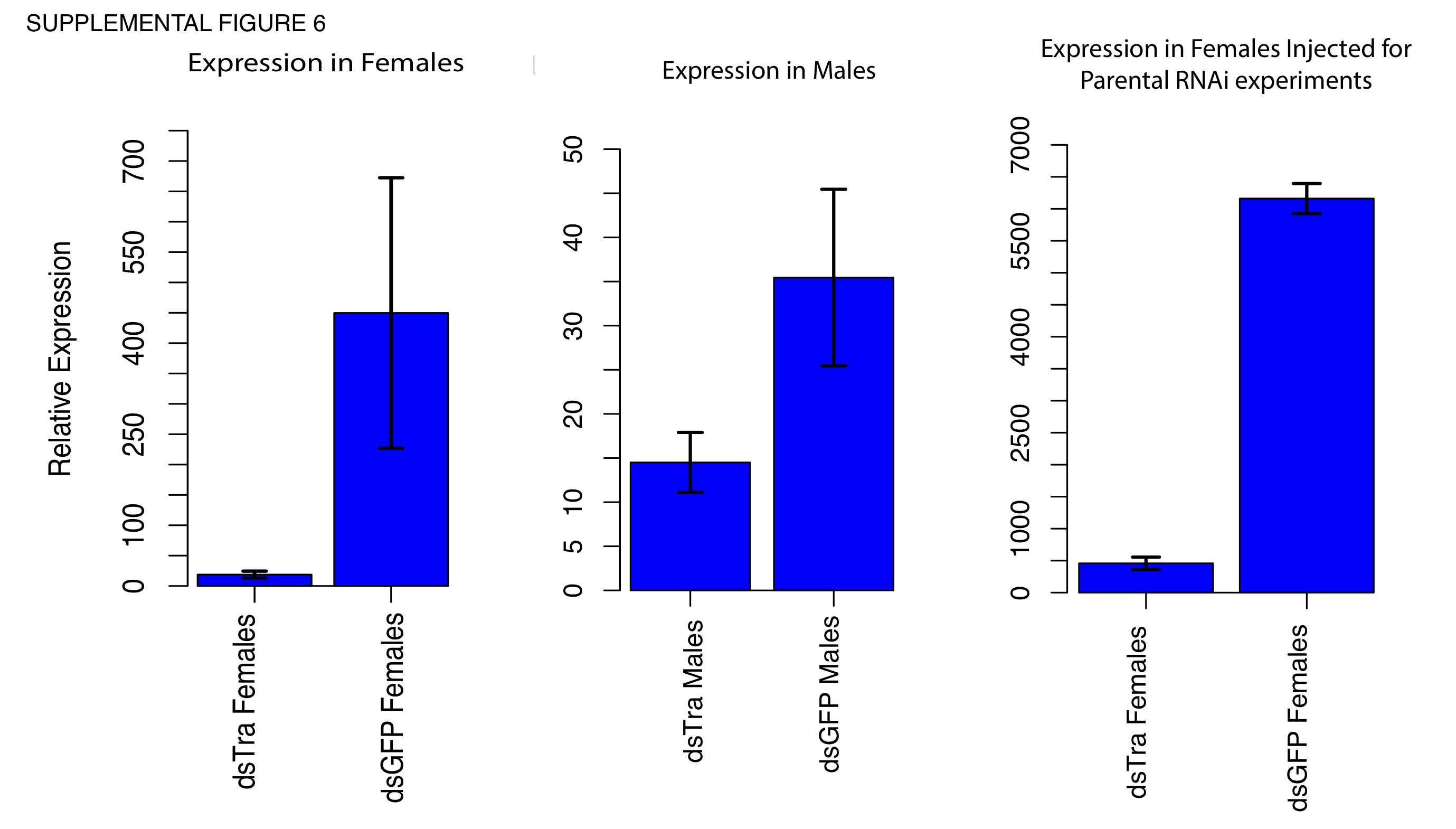

### Supplemental Figure 7

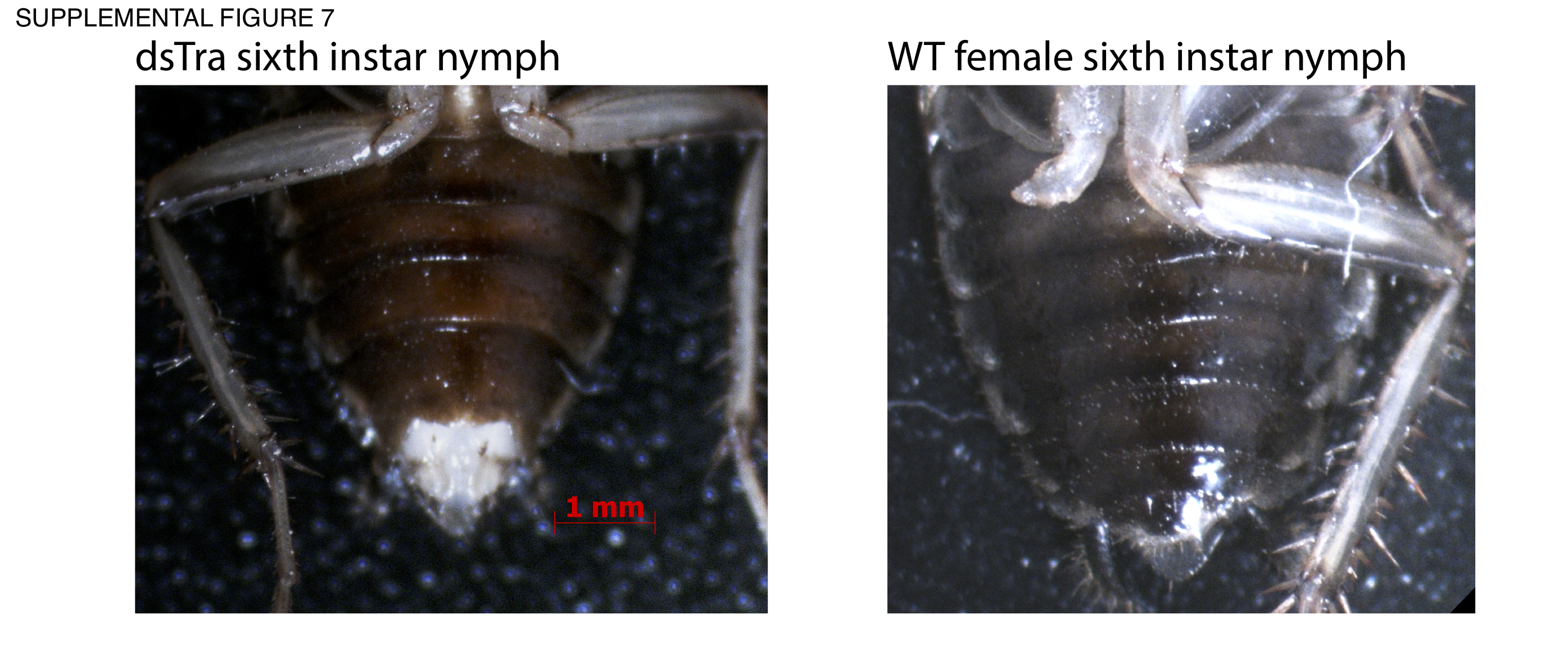

### Supplemental Figure 8

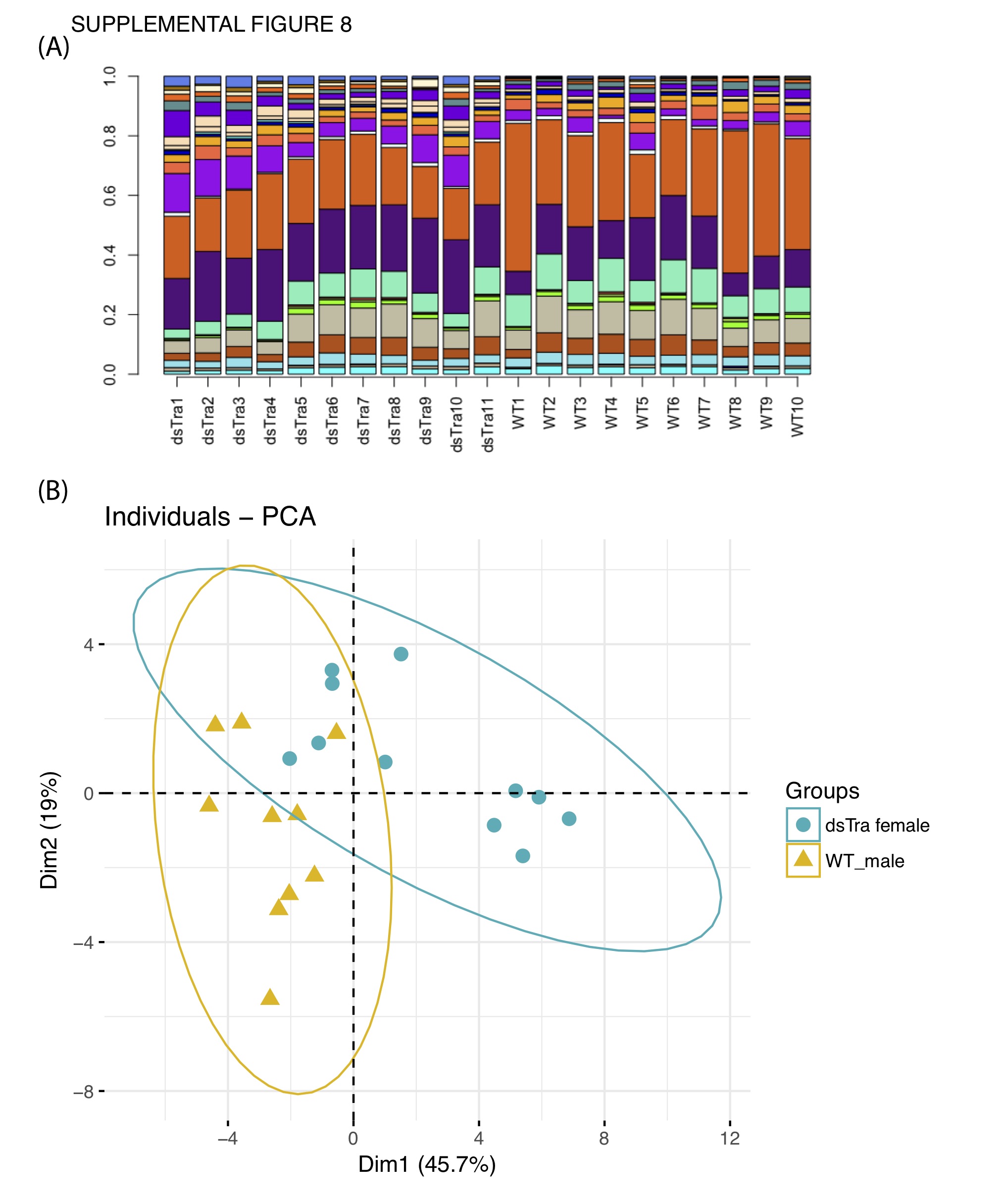

### Supplemental Figure 9

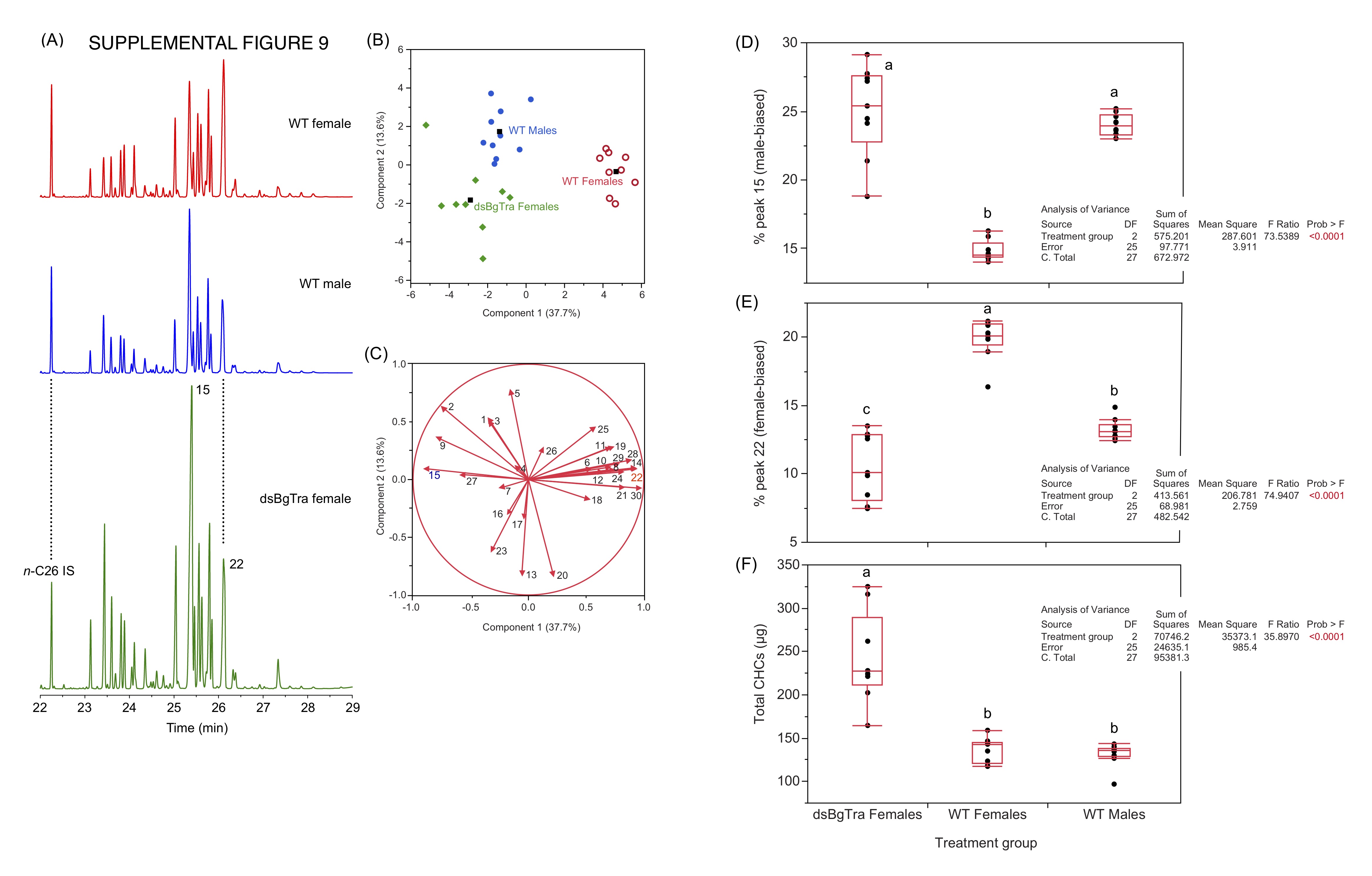

### Supplemental Figure 10

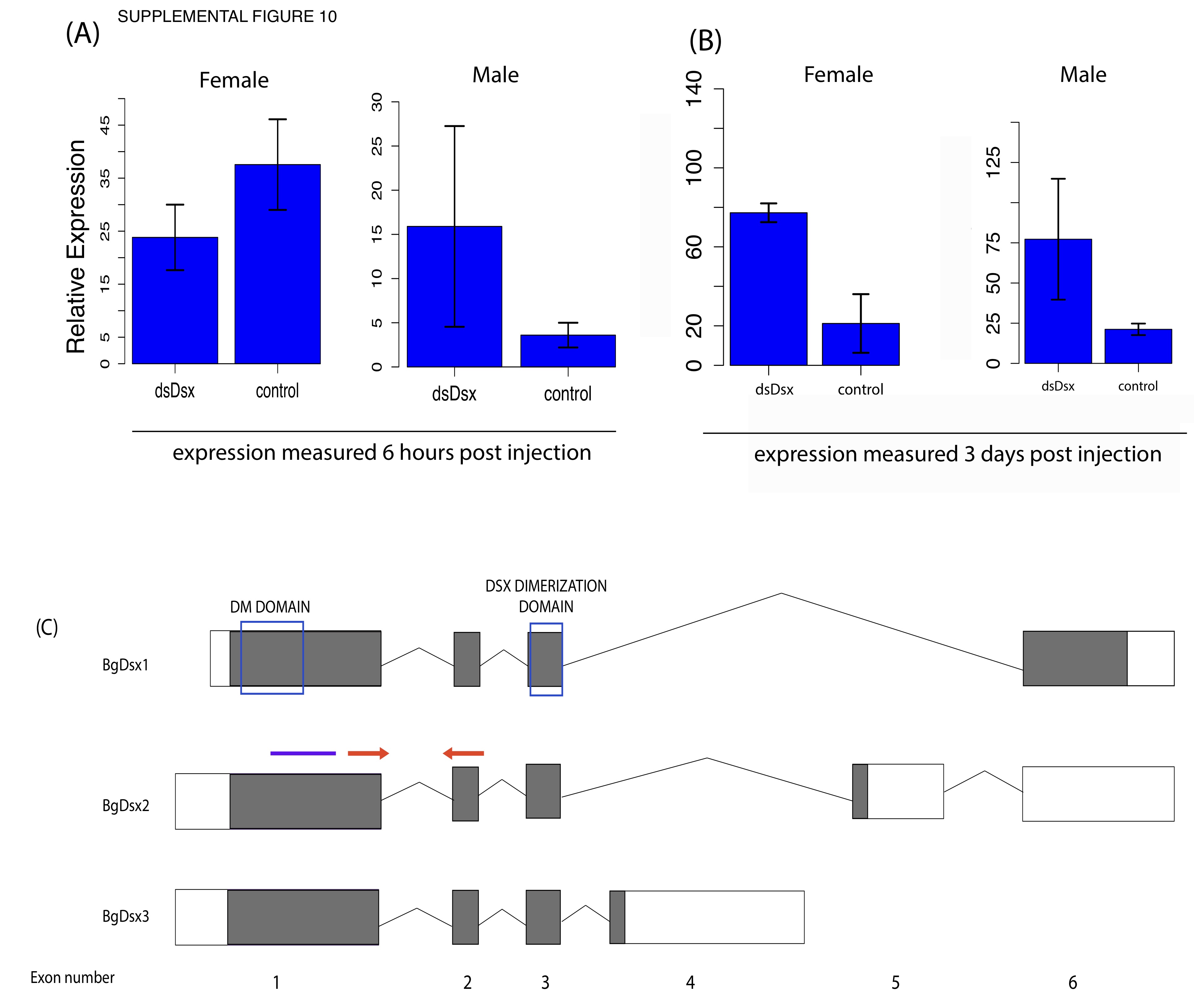

### Supplemental Figure 11

Supplemental Figure 11

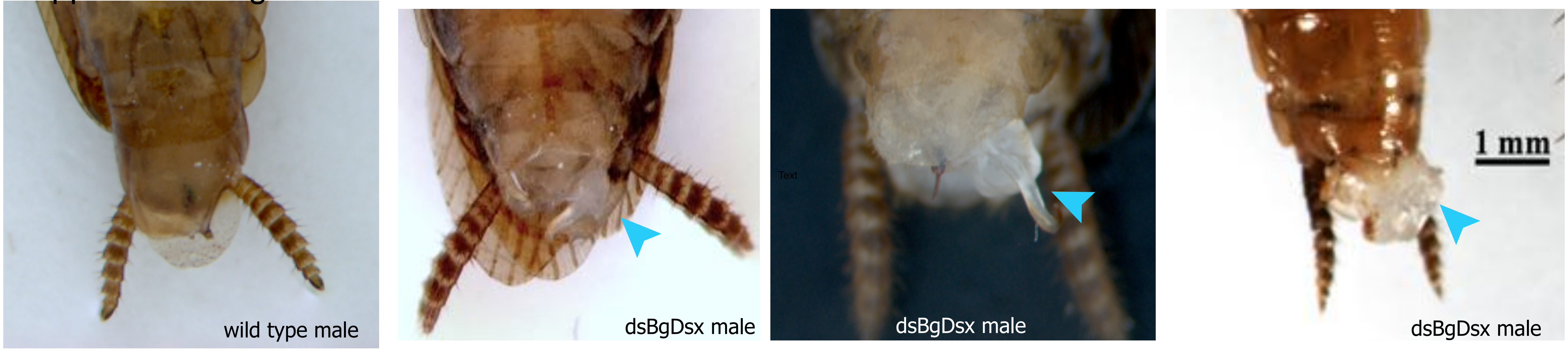

### Supplemental Figure 12

SUPPLEMENTAL FIGURE 12

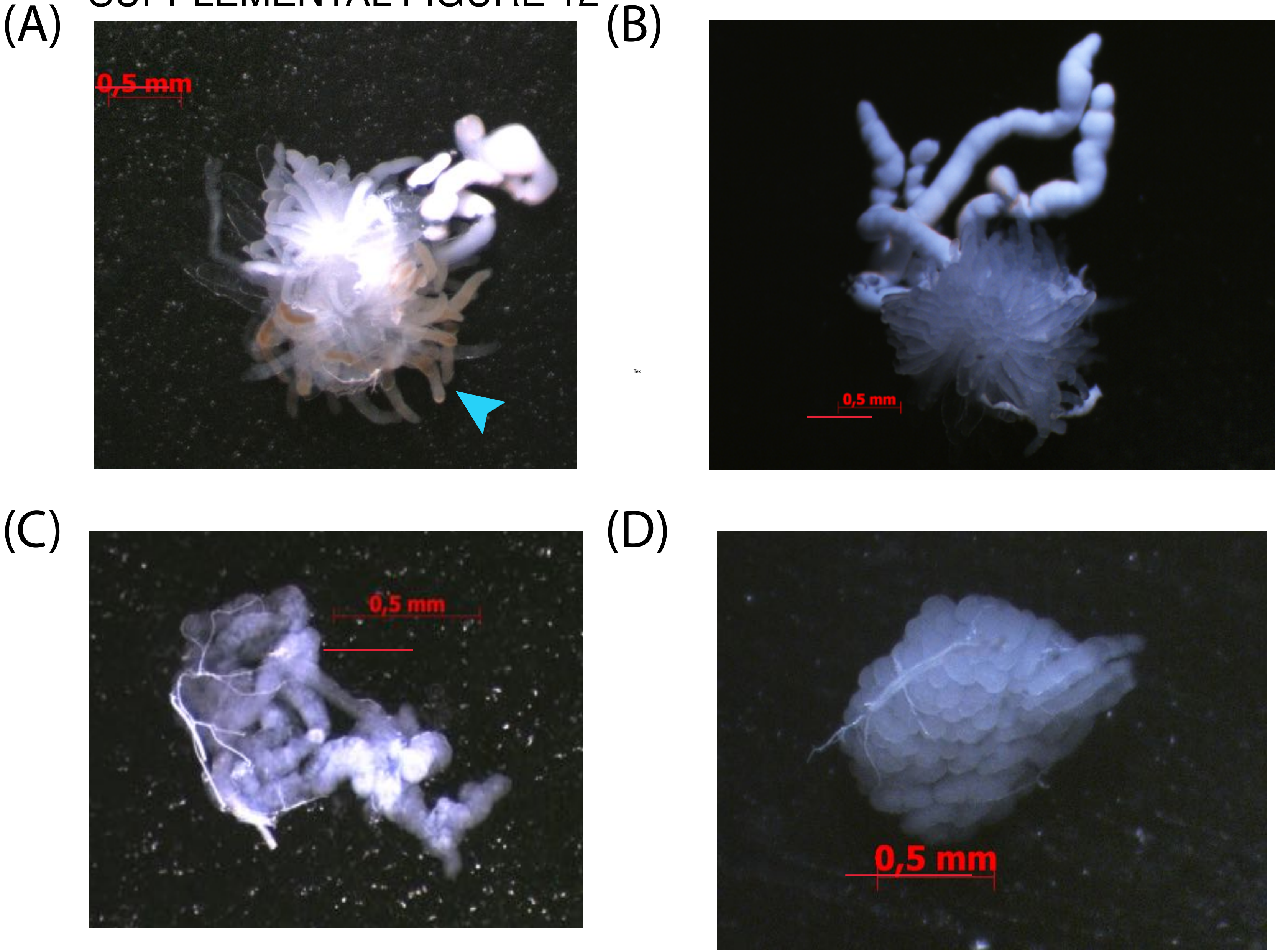
